# Senescent cell-derived extracellular vesicles inhibit cancer recurrence by coordinating immune surveillance

**DOI:** 10.1101/2022.06.30.498366

**Authors:** Tahereh Ziglari, Nicholas L. Calistri, Jennifer M. Finan, Daniel Derrick, Ernesto S. Nakayasu, Meagan C. Burnet, Jennifer E. Kyle, Matthew Hoare, Laura M. Heiser, Ferdinando Pucci

**Affiliations:** Department of Otolaryngology – Head and Neck Surgery, Oregon Health & Science University, Portland, Oregon, US; Department of Cell, Developmental & Cancer Biology, Oregon Health & Science University, Portland, Oregon, US; Department of Biomedical Engineering, School of Medicine, Oregon Health & Science University, Portland, Oregon, US; Biological Sciences Division, Pacific Northwest National Laboratory, Richland, Washington, US; Early Cancer Institute, University of Cambridge, Cambridge, UK; Earle A. Chiles Research Institute, Providence Cancer Institute, Portland, OR, US

## Abstract

Extracellular vesicles (EVs) are key signaling mediators. To explore the role of senescent cell-derived extracellular vesicles (senEVs) in inflammatory responses to senescence, we developed an engraftment-based senescence model in wild-type mice and genetically blocked senEV release in vivo, without significantly affecting soluble mediators. Our results demonstrate that senEVs are both necessary and sufficient to trigger immune-mediated clearance of senescent cells, thereby suppressing tumor growth. In the absence of senEVs, the recruitment of MHC-II+ antigen-presenting cells to the senescence microenvironment was markedly impaired. Blocking senEV release redirected the primary target of senescent cell signaling from antigen-presenting cells to neutrophils.

Through comprehensive transcriptional and proteomic analyses, we identified six ligands specific to senEVs, highlighting their role in promoting antigen-presenting cell–T cell adhesion and synapse formation. Antigen-presenting cells activated CCR2+CD4+ T_H17_ cells, which appeared to inhibit B cell activation. CD4 T cells were essential for preventing tumor recurrence, indicating that CCR2+ T_H17_ cells function downstream of senEVs during senescence surveillance.

Our findings suggest that senEVs complement the activity of secreted inflammatory mediators by recruiting and activating distinct immune cell subsets, thereby enhancing the efficient clearance of senescent cells. These conclusions may have implications not only for tumor recurrence but also for understanding senescence during de novo carcinogenesis. Consequently, this work could inform the development of novel cancer early detection strategies based on the biology of cellular senescence.

## Introduction

Chemotherapeutic regimens such as paclitaxel and cisplatin exert their anti-tumor activity by switching cancer cells to a stable non-proliferative, metabolically active, survival state generally referred to as cellular senescence [1–3]. Senescent cells (SCs) undergo characteristic changes, including an enlarged and flattened morphology, and expression of distinct genes, including senescence-associated β-galactosidase, p16 (CDKN2A), p21 (CDKN1A), [4, 5]. Senescence induction leads to an immediate restriction in cancer cell propagation and tumor growth. However, pathologic accumulation of SCs leads to cancer recurrence [6, 7]. How SCs interact with their microenvironment and the mechanisms that fail to prevent SC accumulation in aging tissues are still largely unknown.

The senescence-associated secretory phenotype (SASP) is a hallmark of senescence [8]. The SASP is composed of several different families of soluble mediators involved in extracellular matrix proteolysis, chemotaxis, cell growth and differentiation, and immune regulation [8–10]. SASP factors are known to both induce paracrine senescence in neighboring cells and to elicit immune responses [11]. Extracellular vesicles (EVs) are important mediators of intercellular communication [12]. SC-derived extracellular vesicles (senEVs) were recently identified as a component of the SC secretome that impact the recruitment of immune cells [13–18]. EVs are membrane-bound, nanometer-sized particles released by every cell type and carrying bioactive signaling molecules from the parental cells [19]. Unfortunately, investigations on the roles of EVs in cancer and senescence have been hampered by the need to isolate them before intravenous reinfusion in animal models; these manipulations can introduce biases such as loss of EV diversity, misjudgment of the amount to reinfuse, and assumptions on EV biodistribution [20]. Model systems that distinguish the contributions of senEVs from those of SASP factors would enable the study the EV functions during immune responses to cellular senescence.

Senescence surveillance is the immune-mediated clearance of SCs. Immune cell types involved in senescence surveillance include monocytes [21, 22], NK cells [23], and T cells [21, 24]. However, SASP-mediated recruitment of other immune cell subsets, such as myeloid-derived suppressor cells [22, 25], can inhibit cytotoxic CD8 T cell responses [3, 6, 7] and promote tumor angiogenesis [24], thereby fostering tumor progression and recurrence. The dual role of the SASP on senescence surveillance may derive from confounding effects due to the presence of different amounts and types of senEVs in SASP studies. Approaches to distinguish between senEVs and SASP would enable to clarify the paradoxical influence of cellular senescence in cancer.

In the present study, we hypothesized that senEVs play a key role in senescence surveillance. To test this hypothesis, we developed and validated an engraftment-based *in vivo* senescence model that minimizes the impact of senescence-inducing stimuli on immune cells while at the same time enabling functional studies of senEVs without impacting the SASP [26–28].

## Results

### A novel engraftment-based senescence model enables study of the senescence microenvironment

Senescent cell-derived extracellular vesicles (senEVs) are a novel and largely underappreciated component of the senescence secretome [29]. To enable *in vivo* assessment of senEVs, we developed an engraftment-based approach that allows us to genetically control EV release specifically in senescent cells (SCs). We induced senescence *in vitro* by treating murine squamous cell carcinoma cell lines (MOC2 and mEER+) with chemotherapeutic agents (Figure 1 and S1, respectively). We chose cisplatin and paclitaxel because they are standard of care for head and neck squamous cell carcinoma (HNSCC) [30, 31]. To optimize the dose, we tested different concentrations of paclitaxel (0-750 nM) and cisplatin (0-35 µM) in a combinatorial manner, with each condition replicated in quadruplicate (Figure 1A). After 48 hours of treatment, we quantified cellular senescence by measuring the senescence marker senescence-associated beta-galactosidase (SA-B-Gal) activity [32]. A defining feature of cellular senescence is the exit from cell cycle, while maintaining cell viability. To confirm that paclitaxel treatment generates *bona fide* SCs, we assessed proliferation (by BrdU incorporation), apoptosis (by Annexin-V) and cell death (by 7-AAD), or lack thereof. These experiments determined the optimal dose of paclitaxel and cisplatin to maximize induction of cellular senescence by minimize proliferation, without inducing significant cell loss by apoptosis or cell death (Figure 1A and S1A). Although maximum senescence was induced when treating MOC2 or mEER+ with 600 nM paclitaxel and 4µM cisplatin, the addition of cisplatin increased cell death and reduced cell recovery, which would have a negative impact on the model. Therefore, cisplatin was excluded from the senescence-inducing treatment in subsequent experiments. Bulk RNA sequencing data from cultured senescent and proliferating cells (MOC2) confirmed increased expression of the senescence markers Cdkn1a (p21), Cdkn2a (p16), Glb1 (SA-B-Gal) and decreased expression of the proliferation marker Ki67 in paclitaxel-treated versus untreated MOC2 (Figure 1B). SCs exhibit an enlarged and flattened morphology [33]. SA-B-Gal activity correlated with morphological changes of the cells after exposure to chemotherapeutic drugs (Figure S1B). These results indicate that 600 nM paclitaxel triggers treatment-induced senescence *in vitro,* while preserving cell recovery.

**Figure 1.**
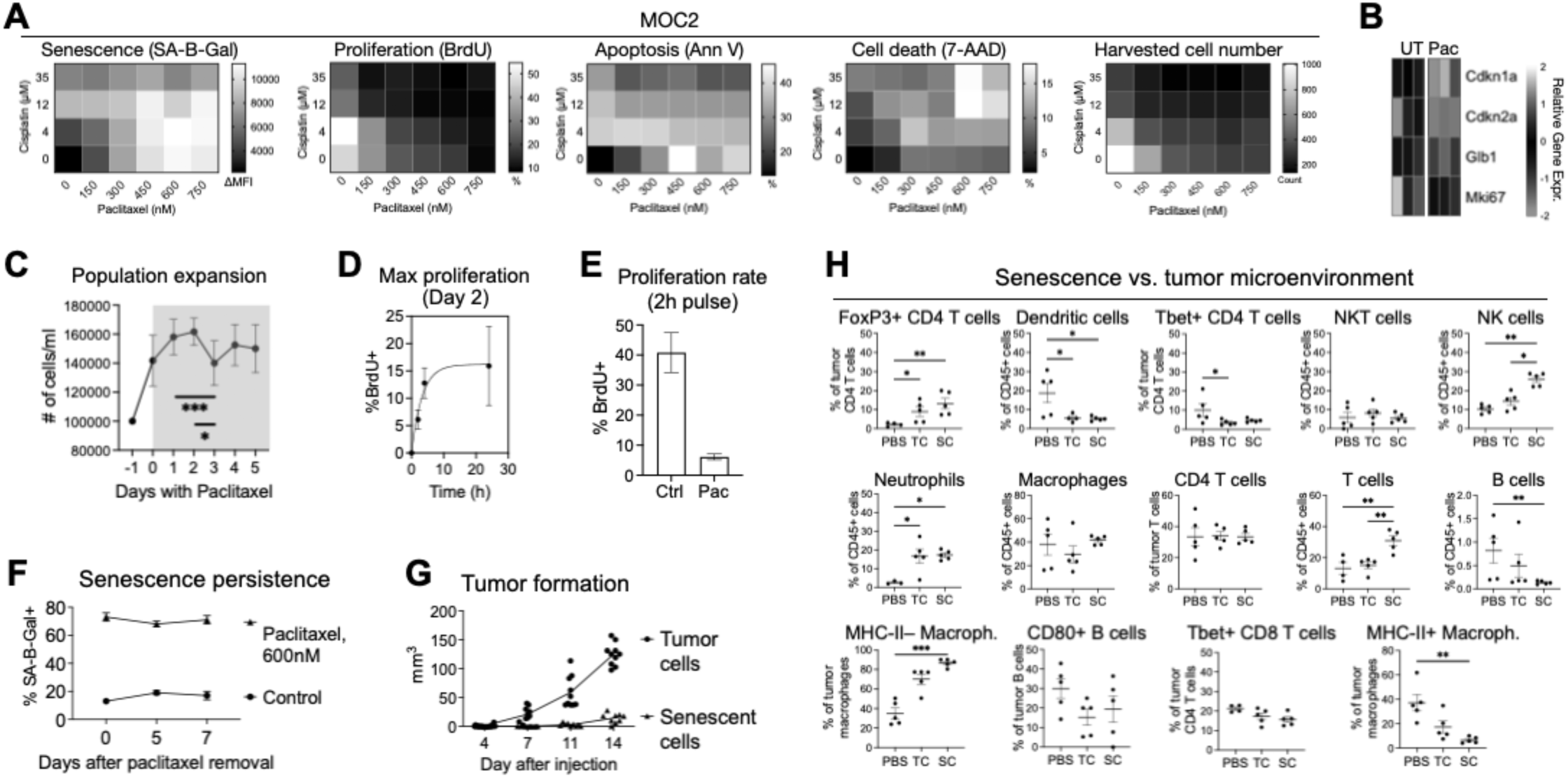
A novel engraftment-based senescence model enables study of the senescence microenvironment. **(A)** Optimization of *in vitro* senescence induction in MOC2 squamous carcinoma cells through 48-hour treatment with specified concentrations of paclitaxel and cisplatin. Ideal conditions ensured elevated senescence levels (quantified by increased SA-β-Gal staining and reduced BrdU incorporation), minimized apoptosis/cell death (measured by annexin V and 7-AAD staining, respectively), and maximization of cell recovery (quantified by cell counting) (n=4). The first column on the left of each heatmap has 0nM paclitaxel and therefore represents cisplatin only. The bottom row of each heatmap has 0uM cisplatin and therefore represents paclitaxel only. The rest of the heatmap shows combination of the two chemotherapeutic agents. **(B)** Differential expression of senescence (Cdkn1a [p21], Cdkn2a [p16], Glb1 [SA-B-Gal]) and proliferation (Ki67) markers in paclitaxel-treated (Pac, 600 nM) vs. untreated (UT) MOC2 cells *in vitro* analyzed by bulk RNA sequencing (n=3). **(C)** Inhibition of senescent cell (MOC2) population expansion by continuous 600 nM paclitaxel treatment for 5 days with fresh media replenishment every 48 hours (n=3, error bars=SD). Statistical analysis conducted using repeated-measure one-way ANOVA with multiple testing correction. **(D)** Inhibitory effect of paclitaxel treatment on MOC2 proliferation over a 24-hour period, resulting in 15% BrdU incorporation (∼7-fold reduction from 100% maximum BrdU incorporation; n=3; error bars=SD). **(E)** Effect of paclitaxel treatment on BrdU incorporation rate in MOC2 cells, demonstrating an 8-fold reduction following a 2-hour BrdU pulse (n=3). **(F)** Persistence of the senescent phenotype *in vitro,* as evidenced by unaltered SA-B-Gal activity for 7 days after paclitaxel removal (n=3). **(G)** Delay in tumor formation by paclitaxel treated senescent MOC2 cells implanted orthotopically in the dermis of mice (1M per mouse, n=10). **(H)** Flow cytometric analysis of immune cell subsets infiltrating tumor or senescent MOC2 cell implants on day 3 post-challenge with 5E5 cells orthotopically (n=5). Statistical analysis conducted using non-parametric one-way ANOVA (Kruskal-Wallis test) with multiple testing correction.

We next quantified the proliferative ability of SCs to evaluate the durability of the paclitaxel-induced senescent phenotype. To this end, we analyzed the *in vitro* proliferation rate of MOC2 cells treated with 600 nM paclitaxel (Figure 1C-E). Continuous paclitaxel treatment blocked cell proliferation, with only a ∼10% cell loss on day 3 as compared to day 1-2 (Figure 1C). To quantify the residual proliferative potential of SCs, we measured BrdU incorporation at saturation and after a short pulse, while culturing SCs in presence of paclitaxel (Figure 1D-E). Over a 24-hour period, BrdU incorporation plateaued at ∼15%, suggesting that paclitaxel alone was sufficient to reduce maximum proliferation of MOC2 ∼7-fold (down from 100% BrdU incorporation). To assess the rate of proliferation, we compared BrdU incorporation between paclitaxel-treated and untreated MOC2 cells after a short pulse with BrdU (2 hours; Figure 1E). We observed an 8-fold reduction in the rate of BrdU incorporation, from 40% to 5%. These analyses indicate that paclitaxel treatment alone can reduce the residual proliferation potential of a fast-growing squamous carcinoma model down ∼2% (see methods).

Since we aimed to implant SCs *in vivo,* without continuing paclitaxel treatment in mice, we asked whether *in vitro* generated SCs retain the senescence phenotype after removal of paclitaxel. To this end, we simulated *in vivo* conditions by removing paclitaxel after two days of treatment and quantified the persistence of senescence over time (Figure 1F). We determined that, over 7 days, SA-B-Gal activity did not decrease and remained present in ∼80% of the cells. These results suggest that the senescent state may be retained *in vivo.* To confirm this, we measured the tumor formation ability of our SC preparation in immunocompetent syngeneic mice (Figure 1G). After implanting the same numbers of viable or senescent MOC2, we observed that SCs were unable to form tumors for at least 2 weeks, indicating that paclitaxel treatment induces a functional state of cellular senescence. These results indicate that our senescence model, while it does not fully recapitulate all the events that take place in patients before and during recurrence (that is, mutagenesis, immune escape, tumor growth, surgery, chemotherapy, recurrence), it can faithfully model the final steps and therefore it represents a useful tool to study tumor recurrence.

The removal of SCs is mediated by the immune system [34]. The immune response against SCs is different from that elicited within the tumor microenvironment [35, 36]. To test whether our engraftment model of senescence recapitulates this difference, we studied the immune infiltrate recruited within 3 days of orthotopic injection of SCs (Figure 1H). When we compared senescence and tumor models, we observed significant differences in their immune composition. As expected [21, 24, 37], the fractions of T and NK cells were significantly increased within senescent tissues 3 days after challenge. Although not statistically significant, we observed a trend in macrophage subsets, with a relative increase in the fraction of MHC-II– CD80+ CD206+ macrophages and a corresponding decrease in the MHC-II+ CD80– CD206– macrophages in presence of SCs. On the contrary, neutrophils, regulatory (FoxP3+) and type-1 helper (Tbet+) T cells were similarly recruited by both tumor and SCs. These results suggest that our senescence model recapitulates the immune responses observed in different *in vivo* models of cellular senescence [21, 24, 37].

### Senescent cell-derived EVs differ from tumor-derived EVs in morphology and protein composition

Extracellular vesicles (EVs) are important mediators of intercellular communication [12]. SC-derived EVs (senEVs) may carry unique signals that impact the recruitment of immune cells ([13–15] and Figure 1H). To explore the role of senEVs in senescence surveillance (that is, the immune-mediated removal of SCs [21]), we characterized the protein composition of senEVs in our model system. We isolated EVs from senescent and proliferating squamous carcinoma cells using ultracentrifugation and compared their protein content, quantities and size using spectrometry, nanoparticle tracking analyses and electron microscopy (Figure 2A-C and S2A). We found that the protein content released into EVs from SCs was ∼6-fold higher compared to non-senescent squamous carcinoma cells (Figure 2A). These results may be explained by either a higher rate of senEV release, larger average senEV size, higher density of protein cargo or a combination of the above. To discriminate between these alternatives, we performed nanoparticle tracking analysis. While EV numbers were similar, we observed a 16% increase in median diameter (Figure 2B), which corresponds to a 56% increase in EV volume (assuming a spherical shape). Electron microscopy analysis confirmed that senEVs are 16% bigger than tumor EVs in both MOC2 and mEER+ cells (Figure 2C). Immunoblot analysis for a pan-EV marker (Flotillin-1; [38–40]) confirmed the presence of *bona fide* EVs (Figure 2D). These findings suggest that senEVs exhibit a larger size and carry a greater protein payload in comparison to EVs from proliferating squamous carcinoma cells.

**Figure 2.**
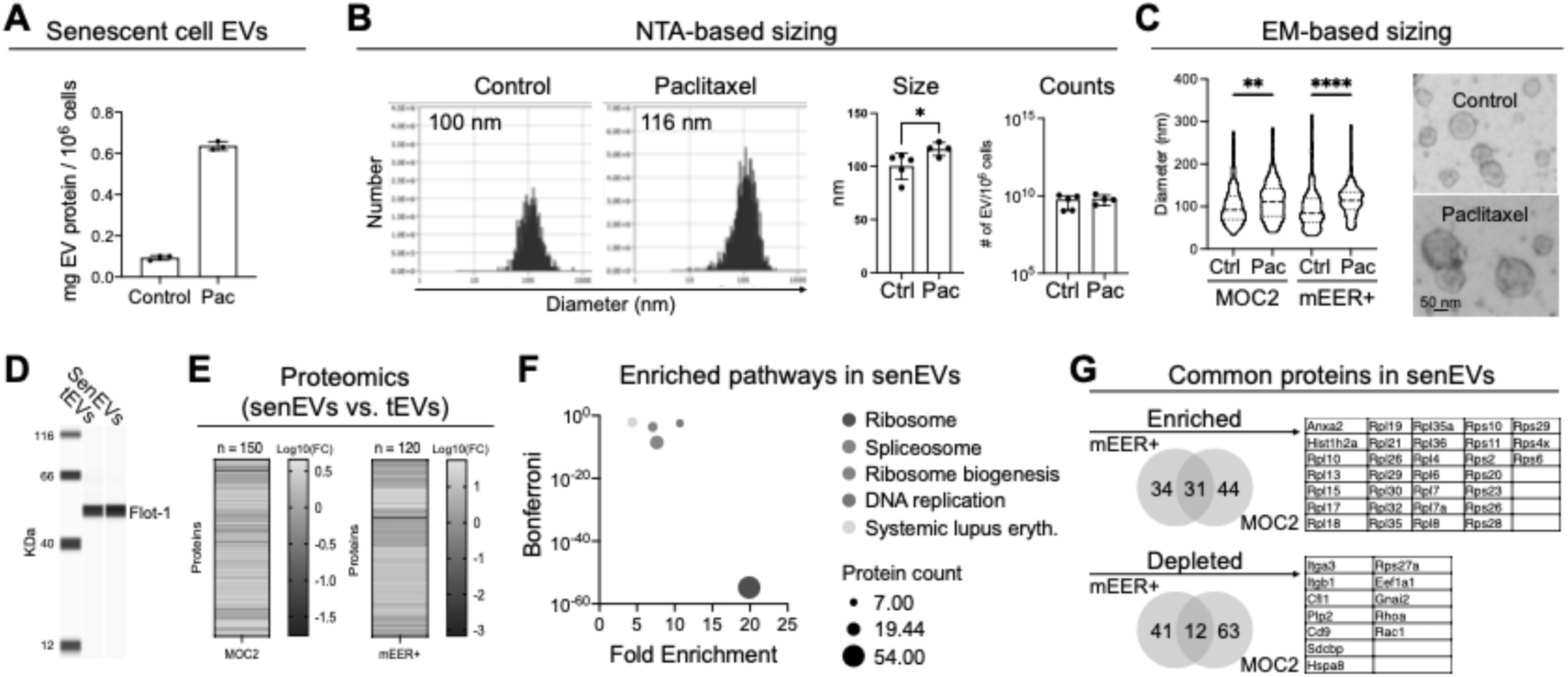
Senescent cell-derived EVs differ from tumor-derived EVs in morphology and protein composition. **(A)** Increased extracellular vesicle (EV) protein content per million senescent MOC2 cells (600 nM paclitaxel for 48 hours), determined to be 6-fold higher compared to non-senescent cells (control), as assessed by NanoDrop (n=3). EV isolation conducted from cell supernatant utilizing ultracentrifugation. Data representative of both MOC2 and mEER+ cell lines. **(B)** Nanoparticle tracking analysis (NTA) of EVs from senescent MOC2 cells: A 16% increase in median diameter corresponding to a 56% increase in volume compared to EVs from proliferating cells (n=5). No differences in EV numbers between proliferating (Ctrl) and senescent (Pac) cell-derived EVs (rightmost graph). **(C)** Electron microscopy (EM) analysis of senescent EVs from both MOC2 and mEER+ cell lines confirms that senEVs are 16% larger than tumor-derived EVs. Data from >10 images per condition. **(D)** Immunoblot for the pan-EV marker Flotillin-1. **(E)** Differential proteomic analysis of EVs isolated from senescent and proliferating squamous carcinoma cells reveals that 150 and 120 proteins exhibit significantly altered expression levels in EVs derived from senescent cells (MOC2 and mEER+, respectively), compared to EVs from proliferating cells (tEVs). **(F)** Pathway analysis (DAVID) of enriched proteins in senescent EVs from senescent MOC2 revealed significant associations with core cellular functions such as ribosomes, spliceosomes, DNA replication, and type-1 immunity (padj<0.01). **(G)** Comparison of proteins enriched in senescent cell-derived EVs (as compared to EVs from proliferating cells) shows a 30% overlap (31 out of 109) between two distinct senescent cell types. Statistical tests were conducted utilizing Mann-Whitney or One-Way ANOVA. Proteomic analysis employed multiple comparisons correction using the Benjamin, Krieger, and Yekkutieli method.

To determine the protein composition of senEVs, we performed proteomic analyses on EVs isolated from senescent and proliferating MOC2 and mEER+ cells (Figure 2E-G). We found 150 and 120 proteins significantly different in senEVs from MOC2 and mEER+ cells, respectively (as compared to EVs from proliferating squamous carcinoma cells, Figure 2E). Pathway analysis showed that proteins enriched in senEVs were significantly associated with core cellular functions, including ribosomes, spliceosome and DNA replication, but also with type-1 immunity (systemic lupus erythematosus; Figure 2F). When we determined the common proteins enriched in senEVs from both cell lines (as compared to EVs from the respective proliferating cells), we found a 30% overlap (Figure 2G). Among these proteins, ribosomal proteins were the largest family.

### Inhibition of EV release does not impact the senescence-associated secretory phenotype

Several studies have reported a dual role for the SASP in both promoting and inhibiting cancer progression and recurrence [3, 6, 7, 21–25]. These conflicting results may derive from the presence of different amounts and types of senEVs in SASP studies. To distinguish the role of senEVs in senescence surveillance from that of the SASP, we aimed to specifically inhibit senEV release. Inhibition of Rab27a is a commonly used method to block EV release [41]. To test if inhibition of Rab27a would stop release of EVs in the context of senescence, we used siRNA oligoes to successfully inhibit expression of Rab27a in senescent cells, as measured by qPCR (Figure S2B). However, two independent methods to assess inhibition of EV release from senescent cells showed that EVs were instead significantly increased (Figure S2C-D). These observations, together with the fact that the total protein content of senEVs was significantly decreased by Rab27a knockdown, suggest that Rab27a in senescent cells may contribute to loading EV cargo proteins and that senescent cells attempt to compensate the diminished EV cargo by releasing higher numbers of senEVs.

We previously validated an alternative genetic tool to inhibit EV release from parental cells [26–28]. In this approach, a dominant-negative mutant of Rab35 (Rab35-DN) is expressed in parental cells. As control, a wild-type Rab35 (Rab35-WT) is used. To confirm that expression of Rab35-DN efficiently and specifically reduces EV release in both proliferating and senescent MOC2 and mEER+, we performed immunoblot assays for the pan-EV marker Flotillin-1 on purified EVs. We observed almost complete absence of EVs in both proliferating and senescent MOC2 expressing Rab35-DN (Figure 3A and S3A). We further validated inhibition of EV release using an independent technique and confirmed a 4-to-6-fold decrease of EV release from SCs expressing Rab35-DN (Figure 3B). Comparison of EV diameters using membrane-intercalating fluorescent dyes (which allows to distinguish membrane-bound particles from non-vesicular particles) suggests that, notwithstanding some variability due to low EV numbers, the increase in size is associated with the senescence phenotype, and not with the expression of Rab35-DN (Figure S3B). These results indicate that the profound inhibition of senEV release is suitable for *in vivo* functional studies.

**Figure 3.**
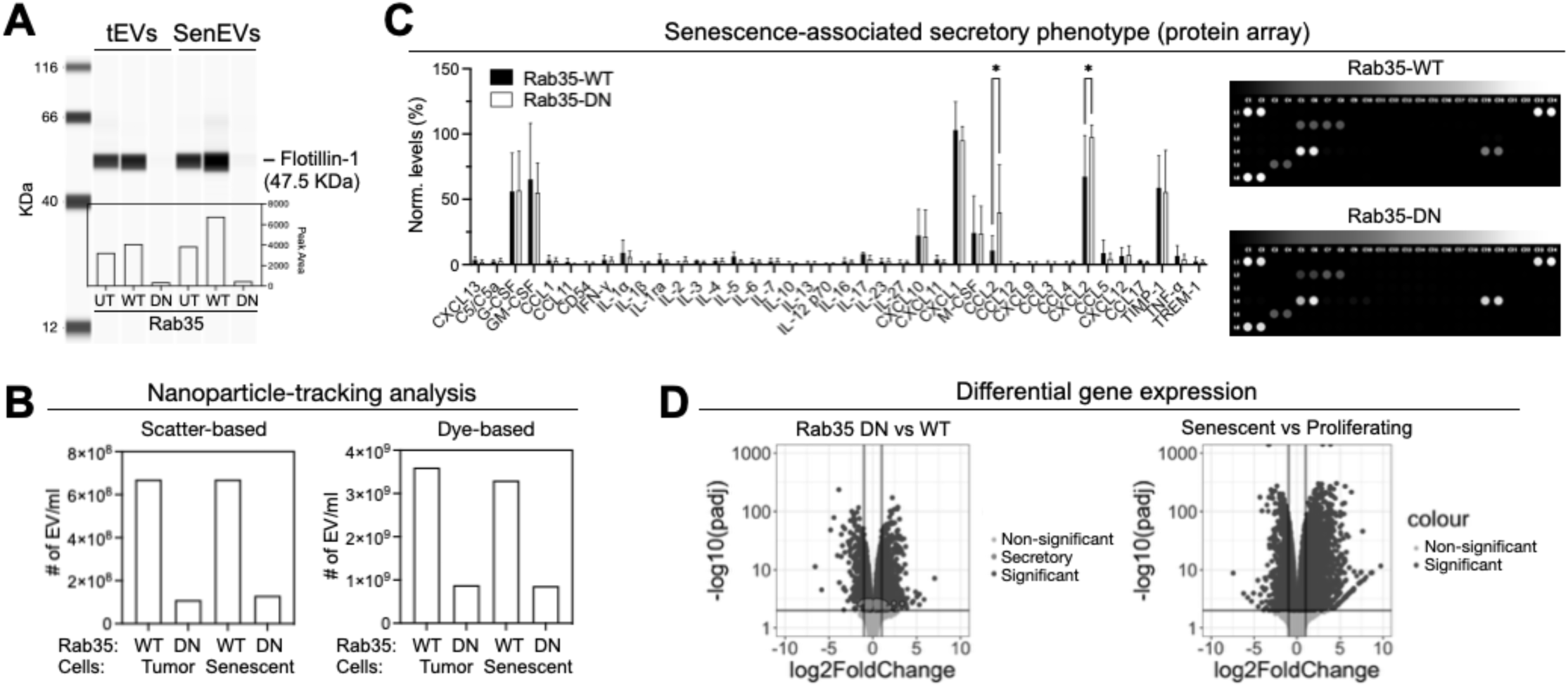
Expression of Rab35-DN does not inhibit the senescence-associated secretory phenotype. **(A)** Effective and targeted reduction of EV release detected in both proliferating and senescent MOC2 cells expressing Rab35-DN using immunoblot assay for the pan-EV marker Flotillin-1. EV preparations were normalized by cell number. UT: untransduced, WT: wild type, DN: dominant negative. **(B)** Detection of reduced extracellular vesicle (EV) release in both proliferating and senescent MOC2 cells expressing Rab35-DN utilizing nanoparticle tracking analysis (NTA), which assesses EV numbers and size via light scatter (left) and fluorescence (right) measurements. **(C)** Senescence-associated secretory phenotype (SASP) factors are not decreased by Rab35-DN expression, as evidenced by quantification of 40 secreted factors using Mouse Cytokine Array on senescent MOC2 cell culture supernatant. **(D)** Bulk RNA sequencing of Rab35-WT and Rab35-DN senescent MOC2 cells (left) and of proliferating and senescent MOC2 cells (right) confirms that Rab35-DN expression has minimal impact on the global SASP (red), defined as genes containing a signal peptide and lacking a transmembrane domain (left); and that changes in gene profile after Rab35-DN expression are minor compared to changes induced by senescence stimuli (right). Significance level set at adjusted p-value < 0.01. Statistical tests by two-way ANOVA with multiple testing correction.

A caveat of interfering with vesicular trafficking via Rab35 is that some SASP factors may be decreased as well, thereby confounding the interpretation of *in vivo* data. To test whether Rab35-DN expression in SCs inhibits SASP components, we quantified 40 different secreted factors by cytokine array on SC culture supernatants (Figure 3C). Of these 40 mediators, 8 were expressed at detectable levels (G-CSF, GM-CSF, CXCL10, CXCL1, M-CSF, CCL2, CXCL2 and TIMP1), and are all well-established SASP factors [9, 42]. Surprisingly, 2 out of these 8 SASP factors (CCL2 and CXCL2) were expressed at higher levels in Rab35-DN SCs. An additional 6 mediators (IL-1α, IL-5, IL-17, CCL5, CXCL12 and TNF-α) were expressed at low but detectable levels and were unaffected by expression of Rab35-DN. To confirm these observations, we performed bulk RNA sequencing of proliferating and senescent MOC2 from cultures (Figure 3D and S3C). We focused our analysis on those genes exclusively featuring signal peptides and lacking transmembrane domains, categorizing them as secretory genes. The analysis of differentially expressed genes between SCs expressing Rab35-DN and SCs expressing Rab35-WT revealed that most secretory genes were not differentially expressed, with only 3 being down-regulated more than 2-fold (Ccl28, Inhbb and Chrd; Figure 3D). Noting that these 3 genes are not commonly found among SASP factors, these findings indicate that expression of Rab35-DN does not impede the release of SASP factors.

### Senescent cell-derived extracellular vesicles recruit antigen-presenting cells and are necessary to inhibit cancer recurrence

While the role of the SASP in inter-cellular communication is well established [9, 10, 43], the specific role of senEVs in communicating with the SC microenvironment is still largely unclear. Although current approaches involving the isolation and injection of exogenous EVs (that is, from cell cultures or biofluids) in animal models permits fine control of pharmacokinetic and pharmacodynamic parameters, it is not clear whether the information obtained from exogenously administered EVs is adequate to address many aspects of EV biology [44]. To overcome these limitations, we took advantage of our engraftment-based senescence model to investigate native senEVs, that is, senEVs released by SCs living within a tissue. To this end, we challenged mice orthotopically with Rab35-WT or Rab35-DN SCs (Figure 4A). We observed that, in absence of senEVs, tumor recurrence was significantly accelerated, as compared to mice receiving Rab35-WT SCs (Figure 4A). To assess whether expression of Rab35-DN interferes with paclitaxel-induced senescence, we compared SA-B-Gal activity and cell expansion *in vitro* (Figure S4A-B). We found that Rab35 manipulation did not influence senescence induction (Figure S4A), and it did not rescue the blockade of cell expansion at the population level (Figure S4B), pointing to the fact that inhibition of senEV release does not interfere with induction of senescence by paclitaxel treatment. To test if inhibition of tumor recurrence by EVs is specific to senescence, we tested if expression of Rab35-DN impacts *in vivo* growth of untreated, non-senescent MOC2 cells. As compared to MOC2 expressing Rab35-WT, we observed no differences in tumor growth of Rab35-DN expressing MOC2 (Figure S4C), indicating that the inhibition of disease progression is a property specific of senEVs. These findings suggest that the release of senEVs inhibits tumor recurrence, possibly via noncell-autonomous mechanisms.

**Figure 4.**
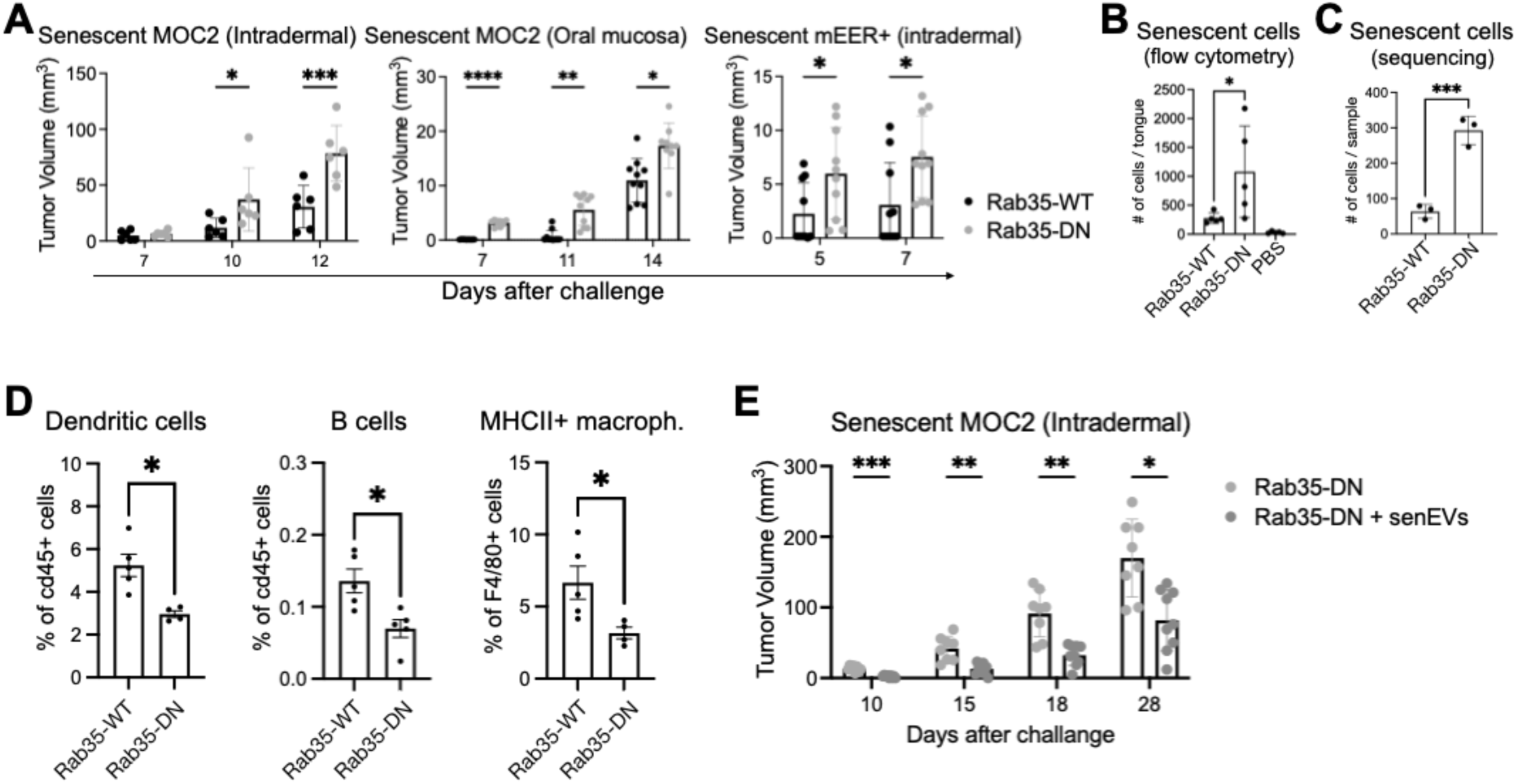
Senescent cell-derived EVs recruit antigen-presenting cells and are necessary and sufficient to inhibit cancer recurrence. **(A)** Rab35-DN expression in senescent cells promotes tumor recurrence in C57BL/6J mice challenged orthotopically (dermis or tongue) with 5E5 senescent cells (three independent experiments, two different senescent cell types, n=10). **(B-C)** Rab35-DN expression in senescent cells promotes accumulation of senescent MOC2 cells 3 days after engraftment, as measured by flow cytometry (B; n=5) and by single-cell sequencing (C; n=3), utilizing the transgenic marker dLNGFR. Data representative of two independent experiments. **(D)** Quantification of immune cell infiltrates by flow cytometry 3 days after implantation of Rab35-WT or Rab35-DN senescent MOC2 cells (data representative of two independent experiments, see Figure S4D). **(E)** Administration of senEVs to mice challenged with Rab35-DN senescent MOC2 cells reverts tumor recurrence (N=10). Statistical tests by one-way ANOVA, two-way ANOVA or Mann-Whitney. Multiple testing correction applied (Sidak).

To better understand the initial host responses to cellular senescence, we studied our engraftment-based model 3 days after challenge with SCs (that is, before tumor formation). We took advantage of a genetic marker (dLNGFR) expressed along with Rab35 variants to track SCs *in vivo* (Figure 4B). We found that interfering with senEV release led to 4-fold higher accumulation of SCs on day 3, as compared to Rab35-WT SCs (Figure 4B). This observation was confirmed by two independent single cell sequencing experiments, also performed on day 3 after SC implantation (Figure 4C). To test if senEVs have a role in SC removal through recruitment of immune cells, we repeated the experiment above and quantified the immune infiltrate early after SC engraftment by flow cytometry (Figure 4D and S4D). Among the immune subsets we queried were CD4 and CD8 T cells, their type-1 effector subsets expressing T-bet, T regulatory cells, NK and NKT cells, myeloid cells (including neutrophils, dendritic cells, macrophages and their subsets), and B cells. We observed that, in presence of senEVs, specific immune cell subsets were significantly increased (∼50%; p<0.05), including infiltrating dendritic cells, B cells and MHC-II+ macrophages (Figure 4D and S4D). These immune cell subsets were related only by the fact that they are all professional antigen presenting cells (APC), indicating that senEVs are necessary to recruit APCs and absence of senEVs increases accumulation of SCs in tissues.

To test whether senEVs are necessary and sufficient to counter tumor recurrence, we performed a complementation experiment in which we reintroduced senEVs along with senescent cells unable to release them. To this end, we implanted senescent MOC2 cells expressing Rab35-DN intradermally in two groups of mice. One group received the senescent cells mixed with 100ug of senEVs, and additional intradermal administrations of senEVs (100ug) every 5 days near the senescent cell injection site. We selected the amount of senEVs based on how many EVs senescent cells release daily *in vitro*. Control mice received the same treatment with PBS (Figure 4E). Strikingly, provision of senEVs restored tumor recurrence to levels observed in untransduced senescent cells (Figure 1G) and in senescent cells expressing Rab35-WT (Figure 4A). Together, these results indicate that the presence of senEVs is sufficient and necessary to counter tumor recurrence.

### Single cell sequencing identifies three distinct subsets of senescent cells *in vivo*

To better understand the signals that senEVs deliver to immune cells during senescence surveillance, we profiled the senescence microenvironment at the single cell level, 3 days after engraftment of Rab35-WT or Rab35-DN SCs, prior to tumor formation. To collect enough cells for profiling, we performed the experiment twice, by pooling 3 mice per group every time. We corrected for differences in sequencing depth using SCTransform [45, 46], which successfully removed batch effects (Figure S5A). We classified three major lineages, namely immune (Ptprc, S100a8, S100a9, Cd14), epithelial (Epcam, Krt5, Krt19, dLNGFR) and stromal (Pdgfra, Pdgfrb, Pecam1; Figure S5B), based on the differentially expressed genes in each lineage compared to the others (Figure 5A). To distinguish between different immune cell subsets, we sub-clustered immune lineage cells separately (Figure 5B). This procedure allowed us to identify a diverse combination of lymphoid and myeloid immune cells recruited to the site of senescence, including two distinct subsets of macrophages characterized by expression of C1q and Spp1, dendritic cells (DC), monocytes, T_H17_ cells, four subsets of neutrophils, and B cells (Table S1-3). The Spp1+ macrophage subset corresponds to the MHC-II+ macrophages identified by flow cytometry (Figure S6A). These results confirmed the presence of the APC subsets we observed by flow cytometric analyses (Figure 4D and S4D).

**Figure 5.**
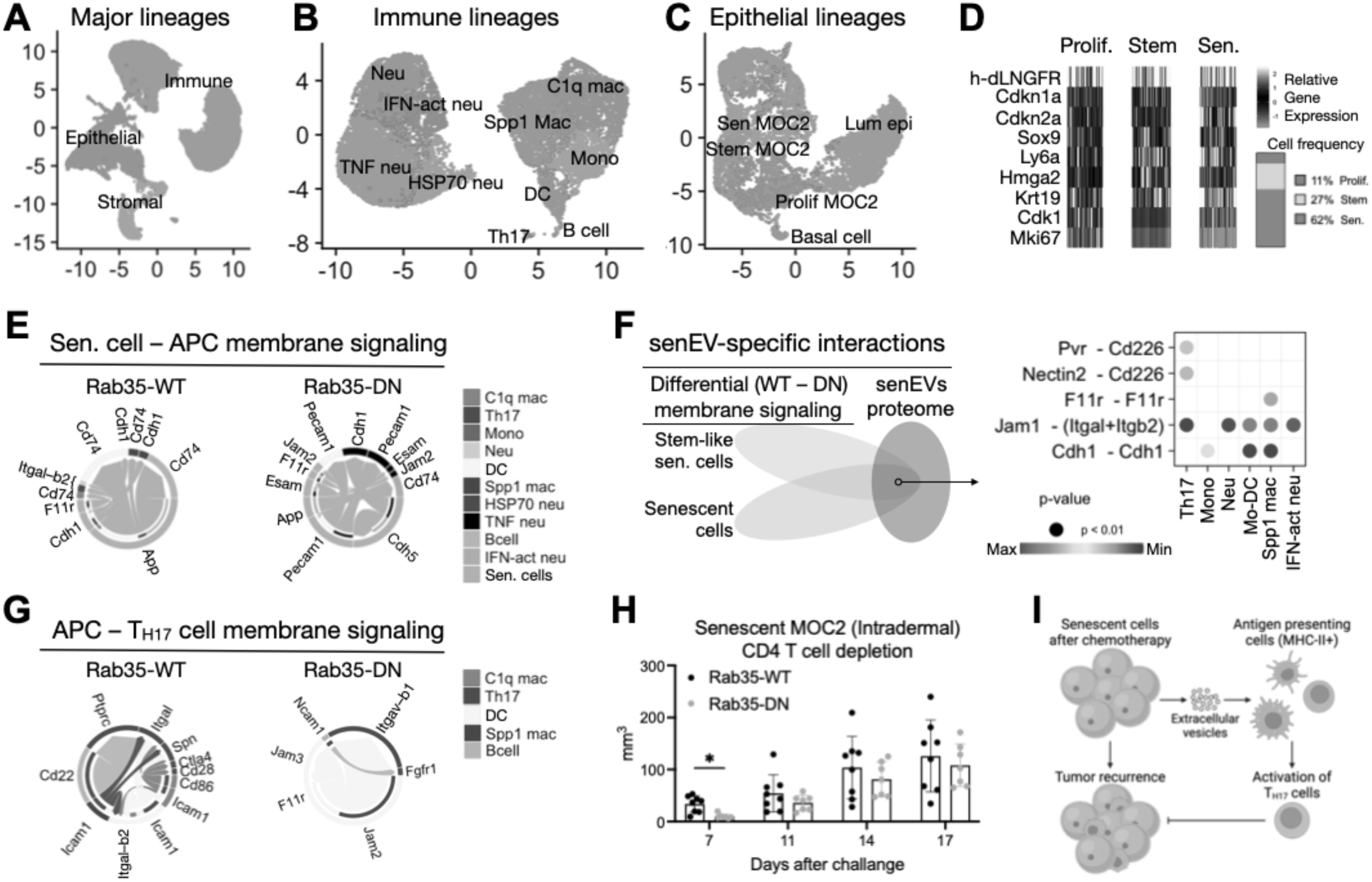
Antigen presenting cells recruited by senEVs require T_H17_ cells to inhibit tumor recurrence. **(A)** Identification of three major cellular lineages within the senescence microenvironment through single-cell sequencing: immune, epithelial, and stromal cells, identified based on differentially expressed genes in each lineage (merged data of two independent experiments). **(B)** Sub-clustering of cells belonging to the immune lineage distinguishes between different immune cell subsets. **(C-D)** Identification of engrafted senescent cell subset among epithelial lineage cells. Proliferating MOC2 cells, stem-like senescent MOC2 cells and senescent MOC2 cells were defined based on markers of proliferation (Mki67), stemness (Sox9, Hmga2, Ly6a/Sca1, and Krt19), and senescence (Cdkn1a and Cdkn2a). **(E)** Visualizing cell-cell communications between senescent cells and immune cells mediated by membrane-bound ligand-receptor interactions using chord diagrams. Each sector in the chord diagrams is an arrow depicting the direction of the ligand-receptor signaling. **(F)** Identification of membrane ligands specifically present on senescent cell-derived EVs. Communications emanating from senescent cells that were enriched in presence of senEVs (differential membrane signaling between Rab35-WT versus Rab35-DN senescent MOC2 cells, left) were intersected with the senEV proteome (right). This analysis was restricted to membrane-bound ligands. The resulting short-list (right) represents putative ligands specifically present on senEVs and not on senescent cell surface. The bubble plot (right) identifies the cellular targets within the senescent microenvironment. **(G)** Visualizing cell-cell communications between antigen presenting cells (APC) and T_H17_ cells mediated by membrane-bound ligand-receptor interactions using chord diagrams. Each sector in the chord diagrams is an arrow depicting the direction of the ligand-receptor signaling. **(H)** Tumor recurrence by Rab35-WT senescent MOC2 cells is increased in absence of CD4 T cells, to levels like those observed in Rab35-DN senescent MOC2 cells. **(I)** Schematic of the proposed mechanism of EVs in senescence surveillance.

To identify the implanted SCs among the single cell pool, we used 3 different approaches: correlation with bulk RNA-seq profiles of SCs from culture, expression of the transgenic marker dLNGFR (co-expressed along with WT or DN Rab35) and copy number variation analysis (Figure S7). This examination identified 3 clusters of epithelial lineage cells as originating from the engrafted SCs (Figure 5C). We named these 3 clusters based on their relative upregulation of markers related to ‘senescence’, ‘stemness’ and ‘proliferation’ (Figure 5D). Notably, all 3 SC clusters were found in both senEV-proficient and -deficient microenvironments, with SCs constituting the largest fraction (62%), followed by SCs expressing the stemness markers Sox9, Hmga2, Ly6a (Sca1) and Krt19 ([47–50]; Figure 5D). These results agree with reports showing that senescent cells can transform into cancer stem cells [51–54], and highlight the usefulness of this model system to further study that process.

### Senescent cell-derived extracellular vesicles display adhesion molecules that can bind to both antigen presenting cells and T cells

The presence of senEVs had a significant impact on which signaling pathways were engaged in immune cells (Figure S6B). To query immune cell–senescent cell communications via EVs, we utilized the R package CellChat to impute likely ligand-receptor signaling from gene expression and distinguish secreted and membrane-bound ligand-receptor interactions [55]. As compared to senEV-deficient, the senEV-proficient senescence microenvironment displayed significant changes in the strength and number of putative ligand-receptor interactions among the 24 cell types (Figure S8A). When expressing Rab35-DN, SCs and stem-like SCs mostly signaled to monocytes, T cells, neutrophils, B cells, luminal epithelial cells and lymphatic endothelial cells (Figure S8A). These results indicate that the senescence microenvironment is highly affected by senEVs.

To focus on membrane-bound ligands (that is, those on the surface of SCs and senEVs), we excluded SASP signaling (see Methods section). When we assessed senescence microenvironments in presence of senEVs, we observed that SCs signaled via membrane-bound ligands mainly with MHC-II+ APCs, that is B cells, DC and Spp1+ macrophages (Figure 5E). In these interactions, the top membrane-bound ligand-receptor pair we identified was App-Cd74, followed by homotypic interactions via Cdh1. Strikingly, when senEVs were absent, the interaction partners of SCs shifted to neutrophil subsets (Figure 5E). SC-Neu interactions in the absence of senEVs were based upon homotypic Pecam1 (CD31) interactions and Cdh5-Cdh1, followed by App-Cd74 and homotypic Esam interactions. Consistently, IFN-activated neutrophils and C1q+ macrophages (that is, MHC-II negative) were both increased in senEV-deficient senescence microenvironments (Figure S8B-C). These results suggest that senEVs promote SC engagement with APCs.

To distinguish between membrane-bound ligands on senEVs versus those on their parental cells (that is, SCs), we took advantage of our senEV proteomic profile (Figure 2E-G). First, we calculated the communications originating from SCs that were enriched in the presence of senEVs. This analysis was restricted to membrane-bound ligands (Figure S9A). Then, we intersected that list with the list of proteins detected on senEVs (Figure 5F). The resulting short-list represented putative ligands specifically present on senEVs and not on SC surface.

This analysis yielded 6 membrane-bound ligands present on senEVs and engaging with cells within the senescence microenvironment: Cdh1, Dsg2, F11r, Ncam1, Ocln, Pvr. These proteins are involved in adhesion and were found on both SCs and stem-like SCs. Interaction analysis suggested that senEVs are capable of binding to both MHC-II+ APC and T_H17_ cells through these adhesion-related proteins (Figure 5F). These results suggest that senEVs may enhance the likelihood of endocytosis of SC antigens or potentially supporting T cell engagement by APCs.

### Antigen presenting cells recruited by senEVs require T_H17_ cells to inhibit tumor recurrence

Given the observed shift in SC-immune cell interactions away from APCs when senEVs are absent (Figure 5E), and the resulting increase in tumor formation (Figure 4A), we hypothesized that adaptive immune responses may have been impacted as well. To test this hypothesis, we queried the signaling pathways from APCs to T_H17_ cells (Figure 5G and S8D). We found that, in senEV-proficient senescence microenvironments, T_H17_ cells have the potential to interact with all three MHC-II+ APC subsets via cell surface signaling (Figure 5G). Moreover, T_H17_ cells expressed the activation marker CD69 (Table S1-3), suggesting that they received antigen-specific stimulation [56]. In senEV-deficient senescence microenvironments, DCs cross talk with T_H17_ cells used different membrane-bound ligand-receptors (Figure 5G), while B cells and Spp1+ macrophages displayed an immunoregulatory role by targeting T_H17_ cells with secreted signals (Figure S8D). These analyses suggest that APCs recruited by senEVs may prime and activate T_H17_ cells.

B-T cell co-activation is central to adaptive immune responses [57]. In senEV-proficient senescence microenvironments, B cell activation was restrained by the inhibitory co-receptors CD22 and Siglecg, and by the release of Galectin-9 (Lgals9) from T_H17_ cells (Figure S8E-F; [58–60]). B-T cell communication via soluble mediators was inferred to be mostly unidirectional, from T_H17_ cells to B cells (Figure S8F). These analyses suggest that T_H17_ cells may inhibit germinal center formation and that B cells may act mainly as APCs during the initial phases of senescence surveillance.

To experimentally validate the role of CD4 T cells in senescence surveillance, we performed antibody-mediated CD4 T cell depletion (Figure S9B). We observed that CD4 T cells were necessary to control tumor recurrence in senEV-proficient senescence microenvironments (Figure 5H). These results are in line with the reported role of this T cell subset in senescence surveillance [21] and suggest that CD4 T cells are downstream of senEVs in the immune response to senescence (Figure 5I).

## Discussion

Murine models have been instrumental in studying cellular senescence across various diseases and physiological processes [61]. They led to the identification of key senescence markers, understanding the role of senescence in disease progression, and exploring potential therapeutic interventions [11]. However, challenges remain in identifying and characterizing senescent cells (SCs) *in vivo* due to the lack of shared markers [62]. Transgenic models of senescence are particularly susceptible to the uncertainty around SC biomarkers and require lengthy characterization of the expression patterns of the gene construct utilized. Non-genetic methods employed to induce senescence in wild-type animals have systemic effects; for example, hydrodynamic (tail vein) injection of large volumes of plasmids expressing oncogenes (such as mutant Ras [21, 22]), or administration of cytotoxic drugs such as Bleomycin [63] cannot control which cell types become senescent. Of note, administration of cytotoxic/cytostatic drugs induces broad damage to tissues and alters immune cell development [64], which complicates the study of immune responses to SCs. Here, we addressed these limitations by validating an engraftment-based senescence model that recapitulates therapy-induced senescence and tumor recurrence in HNSCC patients. While our model system does not fully recapitulate the pathogenesis of HNSCC, we think that our approach specifically models the recurrence stage, in which residual tumor cells accidentally left behind after surgery become senescent *in vivo,* during chemotherapy cycles.

Our approach allows to track the fate of implanted SCs *in vivo* by insertion of a genetic marker (dLNGFR) before senescence induction. The engraftment-based senescence model described here enables tracking of senescent cell fates and functional studies via gene over-expression or knock-down; we took advantage of this feature to inhibit the release of senEVs. Lastly, our model allows us to tightly control which cells are induced into senescence, without affecting immune cells.

The role of EVs in cellular senescence is currently understudied and poorly understood. Some studies reported that senEVs promote tumor cell proliferation *in vitro* [65, 66]. Whether these effects are also in place *in vivo*, and how the senescence microenvironment modulates tumor cell-autonomous effects is unclear. Our results suggest that the noncell-autonomous effects of senEVs may overrule their direct signaling to (pre)-cancerous cells, ultimately establishing senEVs as key to inhibit tumor formation and recurrence. This conceptual advance in our understanding of the senescence microenvironment is in line with the large body of knowledge accumulated around the impact of the tumor microenvironment in cancer [67, 68]. Our results suggest that senEVs are a distinct subset of EVs and differ from them in both morphology and protein content. Importantly, the unique set of proteins carried by senEVs may be involved in recruiting and activating immune cells.

In this study, we engrafted genetically engineered SCs at orthotopic locations to investigate the specific role of senEVs in senescence surveillance, without interfering with the SASP. Although small changes in 5 soluble mediators (Ccl2, Cxcl2, Ccl28, Inhbb and Chrd) were detected after Rab35-DN expression, our complementation experiment indicates that senEVs are sufficient and necessary to counter tumor recurrence. Nonetheless, T_H17_ cells expressed CCR2, which likely mediated their recruitment into the senescence microenvironment [69]. These results suggest that senEVs and SASP factors play complementary but non-overlapping roles in senescence surveillance.

The persistence of SCs in epithelial tissues is detrimental; it is commonly found in precursor lesions leading to HNSCC in patients [70–73], and it can drive cancer recurrence after chemoradiation regimens [6, 7]. Our functional studies indicate that senEVs have a key role in the processes that mediate removal of SCs. This is important because our work shows that accumulation of SCs in absence of senEV signaling leads to increased cancer recurrence. Our analyses suggest that specific immune cells (that is, MHC-II+ APCs) need to be recruited to promptly clear SCs and avoid propagation of the senescence phenotype to neighboring cells. Our data suggest that a potential mechanism for this phenomenon may involve neutrophils, which were increased in the absence of senEVs and have been reported to spread senescence to neighboring cells [74]. On the other hand, APCs like macrophages and dendritic cells, along with IFN signaling, contribute to anti-tumor immunity [75, 76]. Our data shows that APCs release cytokines (including Ccl2, Ccl6 and Ccl7) able to recruit T_H17_ cells via CCR2. Once in the senescence microenvironment, T_H17_ cells signaled with APCs via CD86-CD28, likely leading to T cell activation. Consistently, depletion of CD4 T cells fully abolished senEV protective ability, suggesting that T_H17_ cells are downstream of senEVs in the immune response to senescence. Of note, senEVs carried several cell adhesion molecules that may mediate the uptake of SC antigens by MHC-II+ APCs and may strengthen the immunological synapse between APCs and T cells during senescence surveillance. Overall, these results indicate that senEVs play a major role in senescence surveillance. The ability of SCs to release EVs complements the activity of the SASP in recruiting and activating distinct immune cell subsets for efficient removal of SCs. More work will be necessary to understand if, after uptake, presentation of senEV-bound antigens and generation of antigen-specific immune responses are required to remove SCs.

Virus-like particles (VLPs) from endogenous retrovirus can be found within EV preparations obtained from several mouse tumor cell lines [77]. Our proteomic analysis of senEVs detected high levels of envelop protein from murine leukemia virus, which was present in both senEVs and tumor EVs. Still, it is possible that VLPs may have contributed to the observed effects on tumor recurrence.

Although there may be a small percentage of EVs that are still released by senescent cells expressing Rab35 - DN based on NTA, immunoblot analysis for Flotilin-1, a *bona-fide* pan-EV marker [38–40] suggests a nearly complete (>95%) suppression of EV release. Unfortunately, NTA methods are affected by high background due to the inherent inability of scatter-based measurements to distinguish membrane-bound particles from non-vesicular particles, and the potential presence of micelle-forming dyes during fluorescent-based measurements [78]. We concluded that even if EV secretion was not completely abated, the level of inhibition achieved was suitable for *in vivo* functional studies.

Single cell sequencing data can display significant variability from batch to batch. To account and compensate for this variability, we repeated the single cell analysis twice and performed data transformation, which is a process that removes batch effects. Nonetheless, residual differences may have impacted the results and masked smaller differences.

A limitation of this work is the fact that, in patients, the immune system is also subjected to senescence-inducing stimuli like chemoradiation therapy. It is important to study the interactions between SCs and host cells in stages, by first dissecting how immune cells respond to SCs under normal physiology. Additional studies will compare how senescence-inducing therapies impact these baseline immune reactions [79]. Nonetheless, our conclusions may be directly relevant to understand senescence during *de novo* carcinogenesis and may help devise novel cancer early detection strategies based on the biology of cellular senescence [80].

## Materials and Methods

### Reagents

Paclitaxel from Taxus brevifolia >95% (cat# T7402-1MG) and Cisplatin Pharmaceutical Secondary (cat# PHR1624-200MG), Dulbecco’s Modified Eagle’s (cat# D8062-6X500ML), Dispase II (cat# D4693-1G), BCA protein assay kit (cat# 71285-3), Insulin (cat# I6634-50MG), Hydrocortisone (cat# H0888-10G), Cholera toxin (cat# C8052-.5MG), Transferrin (cat# T1147-100MG), Tri-Iodo-Thyronine (cat# T5516-1mg) were purchased from Sigma-Aldrich). Cellular Senescence detection kit (cat# SG04-10; Dojindo). Amicon ultra-15 centrifugal filters UFC910024 100 kDa (cat# UFC910024), Licor IRDye 800CW Streptavidin (cat # NC0883593), Trypsin EDTA (cat# 25-200-056), RIPA (10X) buffer (cat# 50-195-822) were purchased from Fisher Scientific. Phase-flow Alexa fluor 647 BrdU kit (cat# 370706), BrdU (cat# 423401), APC Anti-BrdU antibody (cat# 364114), Annexin V Binding buffer (cat# 422201), FITC annexin V (cat# 640945), True-Nuclear™ Transcription Factor Buffer Set (cat# 424401), and Zombie Aqua™ Fixable Viability Kit 500 tests (cat# 423102) were purchased from Biolegend. Cell lysis buffer (cat# 9803S; Cell Signaling Technology). Anti-rabbit detection module for Wes, 12-230 kDa Wes Separation module (cat# DM-001) were purchased from ProteinSimple. Proteome Profiler Mouse Cytokine Array Kit, Panel A (cat# ARY006; R&D systems). Penicillin/Streptomycin Solution (cat# SV30010; GE Healthcare). EGF (cat# PHG0311; Gibco). Flow count beads (cat# C36950), Di 8 ANNEPS (cat# D3167), and Pluronic F-127 (cat# P3000MP), Halt Protease Inhibitor Cocktails (PI78430), High-Capacity cDNA Reverse Transcriptase Kit (#4368813), Fast SYBR™ Green Master Mix (#4385616) were purchased from Thermo Fisher Scientific. ultracentrifugation tubes 1 X 3 ½ (cat# NC9146666; Beckman Coulter). Flotillin-1 antibody (cat# NBP2-75492; Novus Biologicals). MACS buffer was prepared by adding 2 mM EDTA (Fisher Scientific), 0.5% BSA (Fisher Scientific) to 500 mL PBS. CD4 depleting antibody (cat# GK1.5; BioXcell). Oligos for RNAi (cat# SR405972; Origene).

### Cell culture

Two oral epithelial squamous cell carcinoma cell lines were used in this study. MOC2 were obtained from Kerafast, Inc., mEER+ were kindly provided by Dr. William Spanos (Sanford Research). MOC2 carries the same mutations observed in human head and neck cancers, namely Trp53, MAPK and FAT [81] whereas mEER+ have been engineered to express Hras^G12V^ and HPV-E6/E7 [82]. MOC2 and mEER+ cell lines were cultured in E-media as previously described [83]. Cell lines were tested for mycoplasma. We prepared E-media by mixing Dulbecco’s Modified Eagle Medium (DMEM) with Ham’s F12 3:1 and supplement with 10% Fetal bovine serum (FBS, from VWR), 0.5% Penicillin/Streptomycin Solution,10 µl of 25 µg/µl Hydrocortisone 16.8 µl of 0.25 µg/µl Cholera toxin, 100 µl of 25 µg/µl Transferrin, 250 µl of 10 µg/µl Insulin, 3.4 µl of 0.2 µg/µl Tri-Iodo-Thyronine, and 250 µl of 10 µg/ml EGF.

### Lentiviral vectors, gene transfer and inhibition of EV release

Lentiviral vector (LV) production and transduction of MOC2 and mEER+ were performed as previously described [26, 84]. Briefly, third generation, self-inactivating LVs were produced by calcium phosphate precipitation. To compare cells expressing Rab35-DN with those expressing Rab35-WT, MOC2 and mEER+ carrying one copy of LV per cell (1CpC) were generated by titration of LVs on cells and purification of transduced cells from the LV dilution that yielded less than 20% positive cells. Purification was performed by magnetic sorting, using dLNGFR, a truncated receptor commonly employed as surface marker [85]. Cells were used as bulk populations, without cloning.

To complement our findings with Rab35-DN expressing cells, Rab27a was targeted using short interfering RNAs (siRNAs). Senescent cells were treated with scrambled control oligos (siScr) or oligos targeting Rab27a (siRab27a) for 48 hours, along with paclitaxel treatment (see below). For oligo transfection, the kit protocol was followed. Briefly, 900 µL of RNAiMAX was diluted with half of the cell media required. Oligos (145ul for siScr oligo and 50ul for siRab27a oligo) were added to 30 mL cell media. The oligo and RNAiMAX dilutions were combined in a 1:1 ratio and incubated for 10 minutes, after which the mixture was added dropwise to the senescent cells in a 15 cm dish. At the endpoint, cell pellets and conditioned media were collected to determine Rab27a gene expression and EV inhibition levels.

### Gene expression

RNA was isolated from senescent cell pellets after siRNA oligo transfection using the Qiagen RNeasy column purification kit and was quantified using a NanoDrop spectrophotometer. Complementary DNA (cDNA) was synthesized from 1000 ng of RNA using the High-Capacity cDNA Reverse Transcriptase Kit. Relative quantification of Rab27a was performed in triplicate with Fast SYBR™ Green Master Mix (Thermo Fischer) using the ΔΔCt method, with GAPDH as reference gene. The following primers were utilized for amplification of Rab27a and GAPDH, respectively (FWD: GAGCAAAGTTTCCTCAATGTCCG, REV: CTTTCACTGCCCTCTGGTCTTC; FWD: CATCACTGCCACCCAGAAGACTG, REV:ATGCCAGTGAGCTTCCCGTTCAG).

### Induction of senescence

To optimize the induction of senescence in MOC2 or mEER+ using chemotherapeutic agents, a matrix of Paclitaxel (0-1000 nM) and/or Cisplatin (0 µM-48 µM) was employed. MOC2 and mEER+ cells were expanded in E-media and plated at 20,000 cells per well in a 96-well plate (in quadruplicate), maintained at 37°C and under 5% CO_2_ for 24 hours. Subsequently, the specified ranges of Paclitaxel and/or Cisplatin were added to the cells, and they were incubated for an additional 48 hours at 37°C in 5% CO_2_. As the stock solutions of paclitaxel and cisplatin were dissolved in DMSO, an equivalent amount of DMSO was added to the control wells.

### Evaluation of senescence induction

#### Senescence-associated beta-galactosidase

The analysis of cellular senescence was conducted using the cellular senescence detection kit SPiDER-B-Gal, and all steps were executed in accordance with the manufacturer’s instructions. Following the 48-hour treatment with Paclitaxel and/or Cisplatin, as previously outlined, the cells were washed with PBS and then incubated at 37°C with 5% CO_2_ for 1 hour after the addition of Bafilomycin A1 solution. Subsequently, the cells were further incubated for an additional 30 minutes after the addition of SPiDER-B-Gal solution. After washing the cells with PBS, they were digested with EDTA-free trypsin, and the resulting cell suspension was collected in a round-bottom 96-well plate containing MACS buffer following centrifugation at 800 rpm for 5 minutes. Finally, a flow cytometer (BD Biosciences) was employed for high-throughput detection, with excitation and emission set at 488 nm and 530 nm, respectively. For flow cytometry analysis, samples that were not treated with spider B gal were utilized to establish the threshold for the negative control.

#### Apoptosis/cell death

To assess the potential unwanted induction of apoptosis and/or cell death by Paclitaxel and/or Cisplatin, parallel plates were prepared. After removing the E-media containing Paclitaxel and/or Cisplatin, cells were washed with PBS, harvested by trypsin, and resuspended in 50 µl FITC Annexin V working solution (3% FITC Annexin V and 3% 7-AAD in Annexin B binding buffer). The cells were then incubated for 15 minutes at room temperature in the dark. Following this, the FITC Annexin V working solution was discarded, and the cells were resuspended in MACS buffer. Quantification was performed using a flow cytometer ( BD Biosciences).

#### Proliferation

Thymidine analog 5-bromo-2-deoxyuridine (BrdU) labeling was conducted following the manufacturer’s guidelines. Briefly, MOC2 and mEER+ squamous cell carcinoma lines were seeded at 20,000 cells per well in a 96-well plate and allowed to reach 70% confluence for 24 hours at 37°C with 5% CO_2_. The following day, cells were exposed to the chemotherapeutic agents and incubated for 48 hours. After washing the cells with media, a pulse of 5 µg BrdU per well was administered for 2, 4, or 24 hours, depending on the study’s objective. Subsequently, cells were washed, fixed, and permeabilized using the corresponding buffers, followed by treatment with DNase for 1 hour. After additional washing steps, cells were stained with 1 µg APC anti-BrdU antibody per well for 30 minutes, washed again, and signals were acquired on a flow cytometer (BD Biosciences). To quantify the residual proliferative potential (RPP) of SCs, we used the following formula:

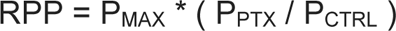

Where P_MAX_ is the fraction of proliferating cells at saturation (that is, after 24h paclitaxel treatment) and P_PTX_ and P_CTRL_ are the short-term (2h) proliferating fractions in paclitaxel or normal media, respectively. Therefore, RPP represents a measure for the senescence-induced reduction in both proliferation rate and maximum proliferation.

### Mice

Six-to eight-week-old female and male C57BL/6J mice weighting between 18 to 25 grams for females and 20 to 30 grams for males were purchased from Charles River Laboratories, housed under conventional SPF conditions and provided with food and water *ad libitum*. Animal care was handled in accordance with the Guide for the Care and Use of Laboratory Animals of the Oregon Health & Science University (OHSU) and covered by OHSU Institutional Animal Care and Use Committee (IACUC). All animal studies, including implantation, tumor size assessment, anesthesia administration, surgery, and euthanasia, were carried out in the OHSU animal facility with a priority on minimizing stress and discomfort. Mice were housed in clean cages with a maximum of 5 mice per cage, furnished with bedding material, a nutritionally balanced diet, and access to clean water ad libitum. The temperature was maintained within the range of 20-26°C, and lighting adhered to a regular light-dark cycle of 12 hours each. Regular visual checks were conducted for health monitoring to promptly detect any signs of distress or illness. Anesthesia was induced using isoflurane and euthanasia was performed in a CO_2_ chamber.

### Engraftment of senescent cells

For tissue engraftments, mice were anesthetized with inhalant isoflurane and the success of anesthesia was confirmed by the lack of reaction to toe pinch. For intradermal senescence model, senescent cells were washed in 50ml PBS to remove residual paclitaxel. A total of 0.5*10^6^ MOC2 or 1*10^6^ mEER+ suspended in 50 µl filtered PBS were injected into flank. For oral senescence model, cells were injected into the central portion of the tongue in a volume of 20 µl. For the intradermal procedure, after removing hair with body trimmer and shaving cream, the beveled tip of a needle with a gauge of 29G or 31G for males or females, respectively, was inserted face up at an angle of less than 10 degrees. The needle was advanced 2-3 mm into the tissue parallel to the skin while simultaneously lifting the needle to create a tenting effect. After injection of the cell suspension, a well-demarcated blister was a sign of successful intradermal placement. Mice in the control groups were injected in the same manner with an equivalent volume of filtered PBS. To track the site of injection for day 3 analyses, a black marker pen was used. Tumor volume was determined by caliper, the greater longitudinal diameter (length) and the greatest transverse diameter (width) were determined twice a week after injection. Tumor volume based on caliper measurement were calculated by the modified ellipsoidal formula as follows:

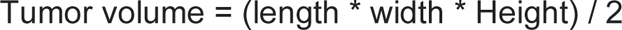

Mice were euthanized by CO_2_ asphyxiation just before tissue harvest. The site of injection was dissected and washed briefly in PBS and weighed. The time points indicated in the graphs as “days after challenge” mean days after injection of senescent cells. Quantification of tumor volumes in the oral mucosa (tongue tumors) was interrupted after two measurements because the mice could not drink and feed themselves anymore. Per IACUC policies, we had to euthanize the cohorts.

### Evaluation of immune response

Samples were dissociated using the following method: tissues were minced with scissors in a 1.5-ml tube in 100 µl dissociation media containing 2% FBS, 1x penicillin/streptomycin, 1x Glutamine, 0.2 mg/ml collagenase IV, 2mg/ml Dispase, and 5U/ml DNase I in IMDM; after transferring to 15 ml conical tubes containing 5 ml dissociation media, minced tissues were incubated at 37 °C with agitation at 500 rpm for 30 minutes in a heater-shaker. After dissociation, the tissues were filtered through a 100 µm strainer with 50 ml of MACS buffer and centrifuged at 500g for 10 minutes. The resulting pellets were transferred to a deep 96-well plate on ice, mixed with pre-diluted surface antibodies and flow count beads. A fraction of the pellets was pooled and stained with corresponding antibodies for fluorescence minus one (FMO) control samples. The plates were sealed with adhesive film and incubated on ice for 30 minutes in the dark. After incubation, the cells were washed twice with PBS and centrifuged at 500g for 5 minutes. Subsequently, they were stained with zombie Aqua dye and incubated for 15 minutes on ice in the dark. Following this, the cells were washed with PBS and gently vortexed to dissociate the pellets. The cells were then fixed and permeabilized using the True-nuclear transcription factor buffer set, following the manufacturer’s guidelines. Briefly, the cells were fixed and incubated at room temperature for 30 minutes. Permeabilization was achieved by adding 1X perm buffer. After three washes in 1X perm buffer, the cells were gently vortexed. A mixture of intracellular antibodies in 1X perm buffer was added to the cells, and they were incubated overnight at 4°C in the dark. Following antibody incubation, the antibodies were removed by washing with perm buffer, and the cells were resuspended in MACS buffer. The readout was performed using a Fortessa flow cytometer.

Flow cytometry analysis of immune cells was according to the following gating:

First *in vivo* experiment:

> Neutrophils: Zombie -, CD45+, Ly6G+, F4/80-
>
> MHC-II+ macrophages: Zombie -, CD45+, Ly6G-, F4/80+, CD80-, MHCII+
>
> Macrophages: Zombie -, CD45+, Ly6G-, F4/80+
>
> B cells: Zombie -, CD45+, Ly6G-, F4/80-, B220+, CD3-
>
> Dendritic cells: Zombie -, CD45+, Ly6G-, F4/80-, B220-, CD3-, MHCII++, NK1.1-
>
> NK cells: Zombie -, CD45+, Ly6G-, F4/80-, B220-, CD3-, MHCII-, NK1.1+
>
> NKT cells: Zombie -, CD45+, Ly6G-, F4/80-, B220-, CD3+, NK1.1+, CD4-
>
> T cells: Zombie -, CD45+, Ly6G-, F4/80-, B220-, CD3+,
>
> T regulatory cells: Zombie -, CD45+, Ly6G-, F4/80-, B220-, CD3+, FoxP3+
>
> T helper 1 cells: Zombie -, CD45+, Ly6G-, F4/80-, B220-, CD3+, CD4+, Tbet+

Second *in vivo* experiment:

> Dendritic cells: Zombie -, CD45+, B220-, F4/80-, CD3-, CD103+, XCR1+, MHCII+
>
> B cells: Zombie -, CD45+, B220+, F4/80-, MHCII+, CD3-, NK1.1-, Ly6G-,
>
> Activated B cells: Zombie -, CD45+, B220+, F4/80-, MHCII+, CD3-, NK1.1-, Ly6G-, CD80+, CD86+
>
> MHC-II+ macrophages: Zombie -, CD45+, B220-, F4/80+, MHCII+, CD206-, CD80-

### Extracellular vesicle purification

Ultrafiltration EV-depleted FBS was prepared as described elsewhere [86]. In brief, regular FBS was subjected to centrifugation in Amicon ultra-15 centrifugal filters for 55 minutes at 3,000 g. The resulting flow-through was collected and used to make E-media. Rab35-WT MOC2 (Rab35 wild-type; capable of EV production), and Rab35-DN MOC2 (Rab35 dominant negative mutant, DN; exhibiting impaired EV production), were cultured in E-media containing EV-depleted FBS and paclitaxel for 72 hours. The absence of EVs in FBS did not impact senescence induction, nor did it change apoptosis or cell death. We used both E-media with and without FBS EVs and we did not notice any difference. Supernatants were collected, normalized based on cell numbers, and then centrifuged at 2000 rpm for 10 minutes to remove cell debris. The supernatant was transferred to ultracentrifugation tubes (Beckman Coulter) and subjected to centrifugation at 24,200 rpm at 20°C for 70 minutes in a SW28 rotor. Following centrifugation, the supernatant was discarded, and 35 ml of sterile, 0.2um filtered PBS was added to the pellet. The mixture was then centrifuged again at 24,200 rpm at 20 °C for 70 minutes. After discarding the supernatants, the tubes were inverted and placed on tissue papers for 1 minute to ensure complete drainage of the tube walls. Subsequently, 50 µl of 0.2µm filtered PBS was added to each tube. The contents were gently pipetted several times to collect EVs, which were then stored at -20 °C until further use.

### Transmission electron microscopy of extracellular vesicles

Electron microscopy of EV preparations was done as previously described [84]. Briefly, 200 mesh-formvar, carbon coated nickel grid were glow-discharged just before use. Five μL of each EV sample (1-5mg/ml of protein) were placed on the grid and incubate for 5 min at RT. After wash 3 x 30ul of filtered PBS (1 min /wash), grids were fixed with 2% glutaraldehyde for 5 min at RT. After washing 5 times with water 2 min each wash, grids were stained with UA 2% 3 min at RT.

### Qualification of extracellular vesicles

#### Western blotting

After successfully purifying extracellular vesicles (EVs) through ultracentrifugation, we proceeded with Western blot analysis using the Simple Western system [87], according to the manufacturer’s guidelines. For this purpose, EVs were lysed using 1X RIPA buffer containing proteinase inhibitor cocktail. The lysate was vortexed several times and mixed with 5X Fluorescent Master Mix. The resulting mixture was then denatured at 95°C for 5 minutes along with a ladder and subsequently transferred to ice. The final concentrations of EV lysate and rabbit anti-Flotillin-1 were optimized at 0.2 mg/ml and 1:50, respectively.

#### Nanoparticle tracking analysis

The quantification of EVs was conducted using nanoparticle tracking analysis (NTA) instrument (Zetaview), employing two independent methods: the first method utilized the properties of both light scattering and Brownian motion to obtain the nanoparticle size distribution of the sample in liquid suspension and the second method evaluated the fluorescence of membrane-intercalating dyes (Di 8 ANNEPS) that have bound to the membrane of EVs. The sensitivity of Zetaview was calibrated to 70 using EV-free media diluted at a ratio of 1:1000 in 0.22um filtered PBS. Subsequently, the number of particles in each sample was determined after diluting the samples at a 1:1000 ratio in filtered PBS with the same calibration adjustment. For fluorescent-based quantification, a Di 8 ANNEPS working solution was prepared by combining 1% Di 8 ANNEPS and 0.25% Pluronic F-127 in PBS. The supernatants containing EVs were then incubated with the Di 8 ANNEPS working solution in a 10:1 ratio for 10 minutes at room temperature. Following the incubation, the samples were diluted at a ratio of 1:1000 in filtered PBS and subsequently applied to the Zetaview, selecting the laser at 488 nm and the filter at 500 nm. Control samples (filtered PBS, filtered media, diluted Di 8 ANNEPS working solution) were routinely used and yielded no particle counts.

### Adoptive transfer of extracellular vesicles from senescent cells

EVs from senescent MOC2 cells were collected as described above. Senescent MOC2 cells expressing Rab35-DN were mixed immediately before injection with senEVs (100ug/mouse, about 25ul) and implanted intradermally as described above (N=10 mice per group). Additional transfers of senEVs (same dosing) were performed every 5 days, intradermally and near the site of senescent cell injection. Control mice received senescent MOC2 cells expressing Rab35-DN mixed with PBS and subsequent PBS injections. The dosing was established based on the amount of senEVs released per day *in vitro.* To avoid batch effects, 3 separate senEV preparation were pooled, each yielding ∼4mg/ml in 250ul. Tumor volumes were measured as described above.

### Mass spectrometry

Proteins were extracted using a modified Folch extraction [88]. Briefly, samples were extracted with chloroform/methanol (2:1, v/v) in a 4-fold excess to the sample volume to yield a hydrophobic layer containing lipids and a protein disc. Each layer was transferred to separate glass autosampler vials and the protein pellet was washed with 1 mL of ice-cold methanol. Each sample was evaporated to dryness *in vacuo*, and appropriately stored at -20°C until further processing and analysis.

Proteins were dissolved in 8 M urea prepared in 50 mM NH_4_HCO_3_, pH 8.0, containing 5 mM dithiothreitol and reduced for 30 min at 37 °C. Cysteine residues were alkylated for 30 min 37 °C with a final concentration of 10 mM iodoacetamide from a 500 mM stock solution. Samples were diluted 8-fold with 50 mM NH_4_HCO_3_, pH 8.0 and 1 M CaCl_2_ was added to a final concentration of 1 mM. Proteins were digested with sequencing-grade modified trypsin (Promega, Madison, WI) overnight at 37 °C. Samples were submitted to solid-phase extraction using C18 spin columns (Ultramicrospin columns, C18, 3- to 30-µg capacity; Nest Group), as previously described [89].

Peptides were separated on a capillary column (60 cm x 50 μm ID packed with C18, 3-μm particles) using a gradient of acetonitrile (mobile phase B) in water (mobile phase A), both supplemented with 0.1% formic acid. The elution was carried with the following gradient: 2 min, 8% B; 20 min, 12% B; 75 min, 30% B; 97 min, 45% B; 100 min, 95% B; 105 min, 95% B; 115 min, 1% B; 120 min, 1% B. Eluting samples were analyzed on a Q-Exactive mass spectrometer (Thermo Fisher Scientific). Full scan spectra were collected in a range of 300 to 1800 m/z with a resolution of 70,000 at m/z 400. The 12 most intense parent ions with multiple charges were submitted to high-energy collision dissociation with 33% normalized collision energy at a resolution of 17,500. Each parent ion was fragmented once and then excluded for 30 sec.

LC-MS/MS data was processed with MaxQuant [90] by searching tandem mass spectra against the human sequences from Uniprot Knowledge Base. Intensity-based absolute quantification (iBAQ) method was used for quantification [91].

### Protein array

The expression levels of cytokines and chemokines in cell supernatants were quantified using the Proteome Profiler Mouse Array Kit. Supernatants from 80% confluent cells were collected and centrifuged at 1000 rpm for 5 min. Array membranes were incubated and developed with supernatants mixed with biotinylated detection antibodies, following the manufacturer’s instructions. For the detection of cytokines/chemokines, IRDye 800CW Streptavidin was used and arrays were scanned using a Li-Cor Odyssey imaging system, which enables near-infrared fluorescence (NIR) detection. The mean pixel intensity for each signal was analyzed using ImageJ.

### Bulk and single cell RNA sequencing

Bulk RNA-seq libraries were prepared using the TruSeq Stranded mRNA Library Prep Kit (Illumina). Briefly, RNA was profiled for integrity on a 2200 Bioanalyzer (Agilent). Poly(A)+ RNA was isolated using oligo-dT magnetic beads and then fragmented using divalent cations and heat. Fragmented RNA was converted to double stranded cDNA using random hexamer priming. The second strand was synthesized with dUTP in place of dTTP to prevent priming from this strand during amplification. The cDNA was ligated to adapters with dual unique indexes and amplified by limited rounds of polymerase chain reaction. The amplified material was cleaned using AMPure XP Beads (Beckman Coulter). The final library was profiled on a 4200 Tapestation (Illumina) and quantified using real time PCR with an NGS Library Quantification Kit (Roche/Kapa Biosystems) on a StepOnePlus Real Time PCR Workstation (Thermo/ABI).

Single-cell RNA-seq libraries were prepared using the Chromium Next GEM Single Cell 3’ Reagent Kit v 3.1 with dual indexes kit (10x Genomics). Cells were counted, then processed through a Chromium Controller (10x Genomics) to produce amplified cDNAs with cell-specific tags. The cDNA was enzymatically fragmented and then size selected using SPRI beads. Adapters were ligated to the cDNA and samples were indexed using indexed amplification primers. Libraries were profiled on a 4200 Tapestation (Agilent) and quantified using real time PCR with an NGS Library Quantification Kit (Roche/Kapa Biosystems) on a StepOnePlus Real Time PCR Workstation (Thermo/ABI). Libraries were sequenced on a NovaSeq 6000 (Illumina) running RTA v3.4.4. BCL files were demultiplexed using bcl2fastq v2.20.0.422 (Illumina).

A custom reference genome was constructed using GRCm38 (Ensembl) and the reporter gene d LNGFR with genome annotations from GRC38.89.gtf. Briefly, the dLNGFR fasta file was added to the end of the GRCm38 fasta as a virtual chromosome using the linux cat command. The gtf file was manually edited to add dLNGFR annotation. The fasta and gtf files were then used to construct a reference genome using the STAR genomeGenerate runMode command. Cell Ranger 7.1.0 uses STAR 2.7.2a aligner for building and aligning to the reference genome. Bulk RNA libraries were aligned using STAR 2.5.0a.

Sequencing data has been deposited to the Gene Expression Omnibus (GEO) under accession numbers GSE262139 (Bulk RNA-seq) and GSE262141 (scRNA-seq).

### Bulk RNA-seq Analysis

Bulk RNA-seq analysis was performed with the DESeq2 (v1.42.0) R package [92]. Raw count data was normalized using the Variance Stabilization Transformation (‘vst’ function) and differential expression was computed between sets of triplicate samples using DESeq2 with LFCshrinkage via apeglm [93].

### scRNA-seq Analysis

All analyses conducted in this study were carried out using R software (version R 4.1.2) and RStudio (Version 2023.12.0+369). Single-cell RNA sequencing data underwent analysis utilizing the Seurat R package (version 4.3) to convert matrix files into Seurat objects [94]. We excluded cells of substandard quality (Minimum number of unique genes = 350 and Maximum Mitochondrial RNA = 15%) . After hashtag demultiplexing (Seurat), doublets were removed and the data from two independent experiments was merged. Both singlets and hashtag negative cells were retained for analysis, with hashtag negative cells used just for dimensionality reduction and unsupervised clustering but excluded from differential expression analysis. UMI counts were normalized with the SCTransform approach to correct for depth of sequencing between cells [95]. The total number of the QC-passed cells in this study was 27,219. The top 2000 highly variable genes were selected for dimensionality reduction using principal component analysis (PCA) and uniform manifold approximation and projection (UMAP). Initial cell clustering was performed with the top N principal components using the Louvian algorithm (‘FindClusters’ function) across a sweep of resolutions. We computed the silhouette width and Root Mean Squared Error (RMSE) for each clustering resolution and used the value that maximized silhouette width and minimized RMSE [94]. The “FindAllMarkers” function was utilized to identify marker genes upregulated in each cluster and define their identity, resulting in the identification of 23 clusters, including 7 stromal clusters, 5 epithelial clusters, and 9 immune clusters. This approach failed to resolve dendritic cells and B cells, so the cluster containing both was separated and reprocessed (variable feature identification and PCA) to identify them, increasing the total number of clusters to 24. Additionally, the Seurat function ’FindAllMarkers’ was used to identify differentially expressed genes (DEGs) for each cluster.”

We used three metrics to differentiate between epithelial clusters composed of endogenous cells and the implanted MOC2 SCs; 1) Imputed Copy Number Variation distance from diploid, 2) Correlation to bulk MOC2 RNA-seq profiles, 3) Expression of human transgene dLNGFR (Supplemental Figure 7). For copy number variation analysis, we subset the scRNA-seq data to just the cells with either a ‘stromal’ or ‘epithelial’ lineage label and used the CopyKAT R package (v1.1.0) to infer Copy Number Variations from gene expression [96]. The CopyKat algorithm was applied with ‘stromal’ cells set as expected diploid using the ‘mm10’ genome and default parameters. To identify outlier cells, we computed the distance from diploid as the logarithm of summed squared distance from diploid. To compute expression correlation to bulk profile, we found the intersect of genes expressed in the scRNA-seq and bulk-RNAseq data and computed the single cell Pearson correlation to mean expression bulk profiles for triplicate samples of paclitaxel treated MOC2-25 (EV Proficient) and MOC2-27 (EV Deficient). For assessment of dLNGFR expression, this gene was added to our reference transcriptome pre-alignment and dLNGFR was normalized with the SCTransform approach along with all endogenous mouse genes.

For intercellular communication analysis and network visualization of scRNA-seq data, we used the R package “CellChat” (v1.6.1), a public repository of ligands, receptors, cofactors, and their interactions [55]. We utilized CellChatDB.mouse, a database containing literature-supported ligand-receptor interactions in mice. This database distinguishes between three classes of ligand-receptor interactions: secreted signaling, ECM-receptor, and cell-cell contact interactions. Seurat normalized gene expression was used to create cell chat objects from the pooled tumors for each condition (senEV proficient, senEV deficient), and then these interactions were compared to identify differences across the conditions. To identify signaling pathways between senEVs and their microenvironment, we focused our analysis specifically on membrane-bound ligand-receptor interactions (that is, cell-cell contact interactions). A heatmap was used to detail the differential interaction numbers and strengths between the different subsets identified within the senescence microenvironment. Chord diagrams were employed to focus on top membrane-bound ligand-receptor interactions, after specifying the sender and recipients of interest. A bubble plot was used to compare communication probabilities mediated by specific ligand-receptor pairs (based on proteomic data) toward any immune cell subset. SCs and stem-like SCs displayed overlapping interaction partners, including themselves, IFN-activated neutrophils, basal epithelial cells and Schwann cells, when expressing Rab35-WT. For this reason, we considered them as a single subset in following analyses.

To unbiasedly define SASP factors, we filtered BioMart database [97] for genes containing a signal peptide and lacking a transmembrane domain.

### CD4 T cell depletion

To deplete CD4 T cells, 200ug of anti-CD4 antibody (GK1.5) was administered intraperitoneally once per week. The first administration was performed a day before challenge with senescent MOC2 cells. More than 99% of CD4 T cells were depleted in circulation as confirmed 4 days after the first GK1.5 treatment (using a different anti-CD4 clone).

### Statistics

Statistical analyses of the results were performed using appropriate tests in GraphPad Prism version 10. Proteomic and RNA sequencing data were analyzed using R. Adjusted P values < 0.05 were defined as significant. Biological and technical replicates are reported for each experiment.

## Acknowledgements

We thank Dr. Robert Searles, Director of the Massively Parallel Sequencing Shared Resource at OHSU, for help and advice with sequencing and genome alignment. We thank the OHSU Multiscale Microscopy Core for help with electron microscopy. This work was funded by the department of Otolaryngology startup grant (FP), CEDAR-CRUK award CRUK6851219 (FP, MH), NIH T32 Fellowship in “Integrated Training in Quantitative and Experimental Cancer Systems Biology” CA254888 (TZ), and by Exploratory Research Seed Grant funding from the OHSU School of Medicine. This research is affiliated with the PMedIC joint research collaboration between OHSU and PNNL. The proteomics mass spectrometry analysis was performed at the Environmental Molecular Sciences Laboratory a U.S. Department of Energy National Scientific User Facility located at the Pacific Northwest National Laboratory operated under contract DE-AC05-76RL01830.

