## Supplementary figures for "Senescent cell-derived extracellular vesicles inhibit cancer recurrence by coordinating immune surveillance"

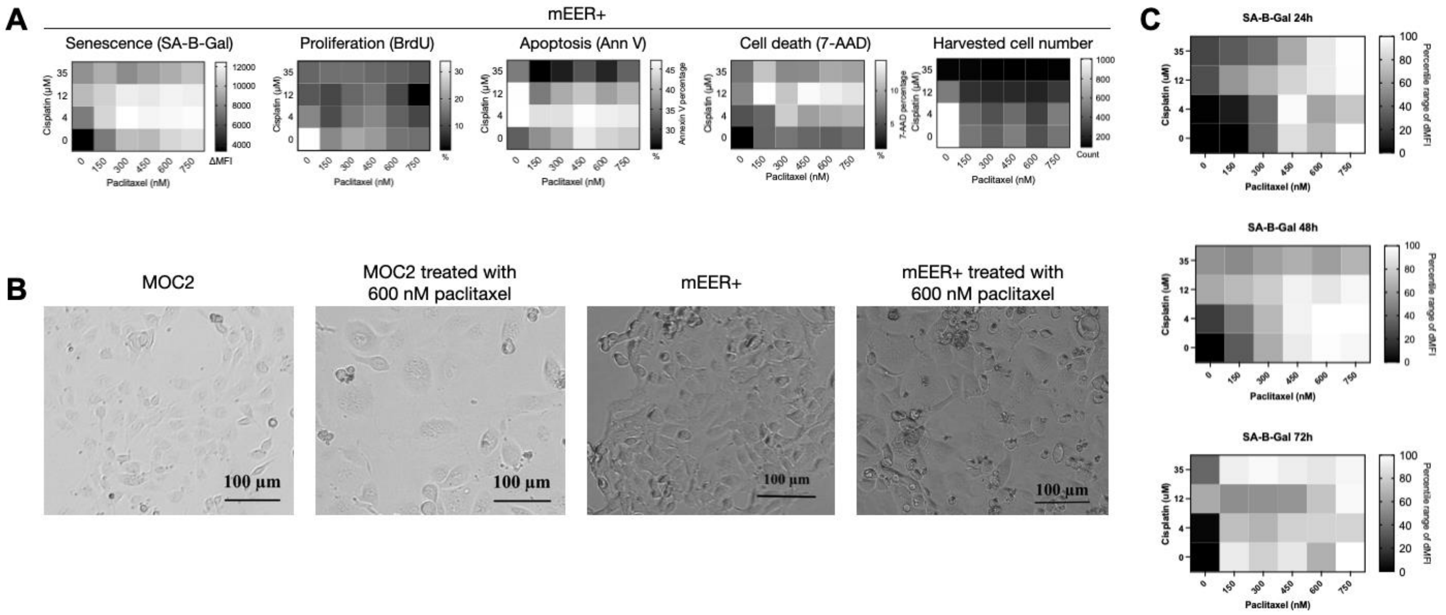


#### Figure S1. Validation of *in vitro* senescence induction using chemotherapeutic agents.

**(A)** Optimization of senescence induction in mEER+ squamous carcinoma cells through 48-hour treatment with specified concentrations of paclitaxel and cisplatin. Ideal conditions ensured elevated senescence levels (quantified by increased SA-β-Gal staining and reduced BrdU incorporation), minimized apoptosis/cell death (measured by annexin V and 7-AAD staining, respectively), and maximization of cell recovery (quantified by cell counting) (n=4). **(B)** Microscopy images revealing enlarged and flattened morphology of MOC2 and mEER+ cells following treatment with 600 nM paclitaxel for 48 hours. **(C)** Determination of ideal duration of paclitaxel treatment by SA-β-Gal staining. MOC2 cells were treated for 24h, 48h or 72h with 600 nM paclitaxel. Normalized data is reported.

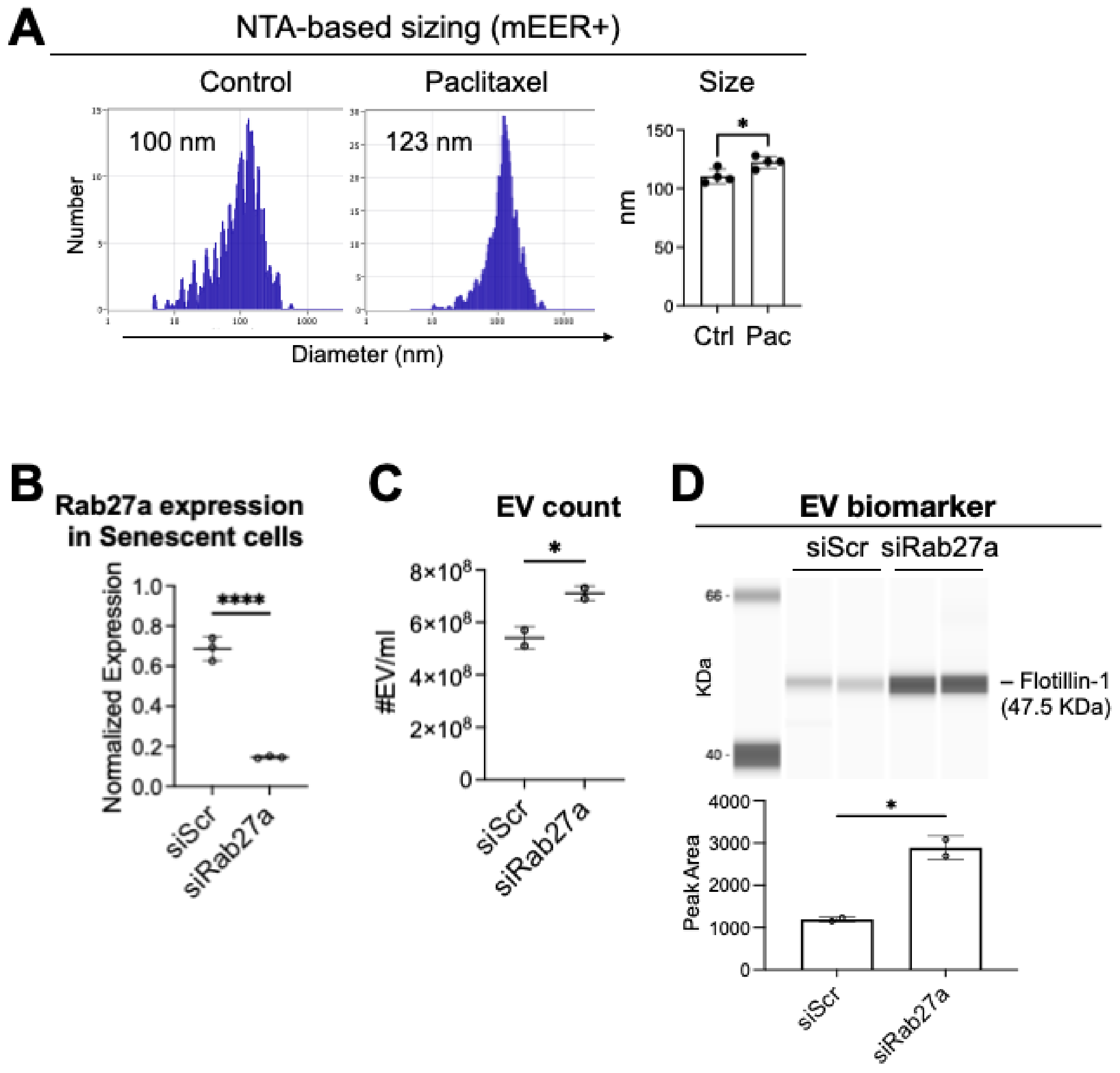


Figure S2. Rab27a inhibition decreases senEV content and increases their numbers

**(A)** EVs from senescent and proliferating mEER+ cells were sized by nanoparticle tracking analysis (NTA). Similarly to MOC2, also mEER+ senEVs are bigger than mEER+ tEVs. **(B)** Gene expression quantification (qPCR) for Rab27a after RNA interference in senescent MOC2 cells show strong suppression of Rab27a mRNA. **(C)** NTA quantification of senEVs from senescent MOC2 cells transfected with either a scrambled siRNA oligo (siScr) or an siRNA oligo targeting Rab27a (siRab27a) shows that Rab27a inhibition increases the concentration of senEVs in senescent MOC2 media. Numbers are normalized by senescent cell counts. **(D)** Immunoblot for the pan-EV marker flotillin-1 on samples from (C) confirms that Rab27a inhibition increases the amount of senEVs in senescent MOC2 media. Numbers are normalized by senescent cell counts. **(E)** Spectrophotometric quantification of total protein content on samples from (C), normalized by senescent cell counts, shows that Rab27a inhibition decreases cargo loading.

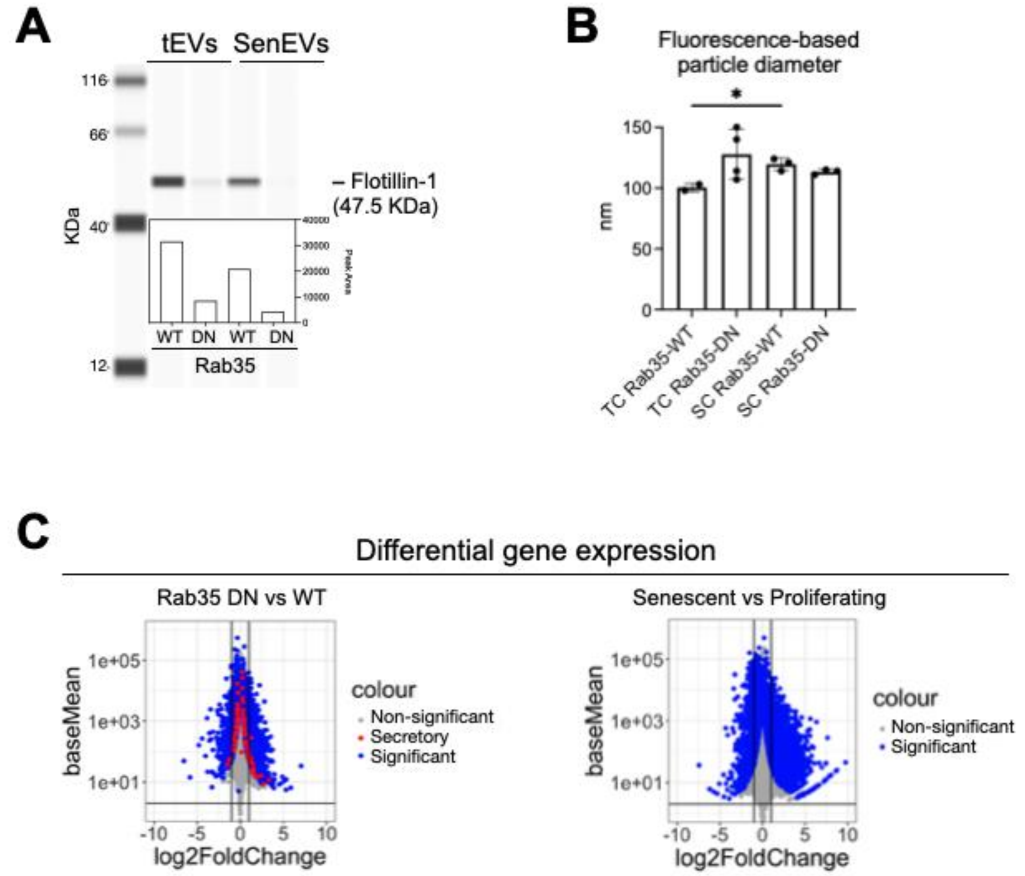


Figure S3. Expression of Rab35-DN does not inhibit the senescence-associated secretory phenotype

**(A)** Immunoblot analysis of EV preparations from mEER+ cells expressing either Rab35-WT or Rab35-DN. Senescence was induced with 600nM paclitaxel. The pan-EV marker Flotillin-1 was assayed. **(B)** NTA quantification of EV diameters using membrane-intercalating fluorescent dyes (which allow to distinguish membrane-bound particles from non-vesicular particles) indicates that the increase in size is associated with the senescence phenotype (compare group 1 vs 3), and not with the expression of Rab35-DN (compare group 3 vs 4). This is consistent with data in Figure 2B-C. **(C)** Bulk RNA sequencing of EV-proficient and EV-deficient senescent cells (left) and of proliferating and senescent MOC2 cells (right) confirms that inhibition of senEV release has minimal impact on SASP. Secretory genes (red) were defined as those containing a signal peptide and lacking transmembrane domain (left). Significance level set at adjusted p-value < 0.01. Expression level is indicated by “baseMean” (related to Figure 3D).

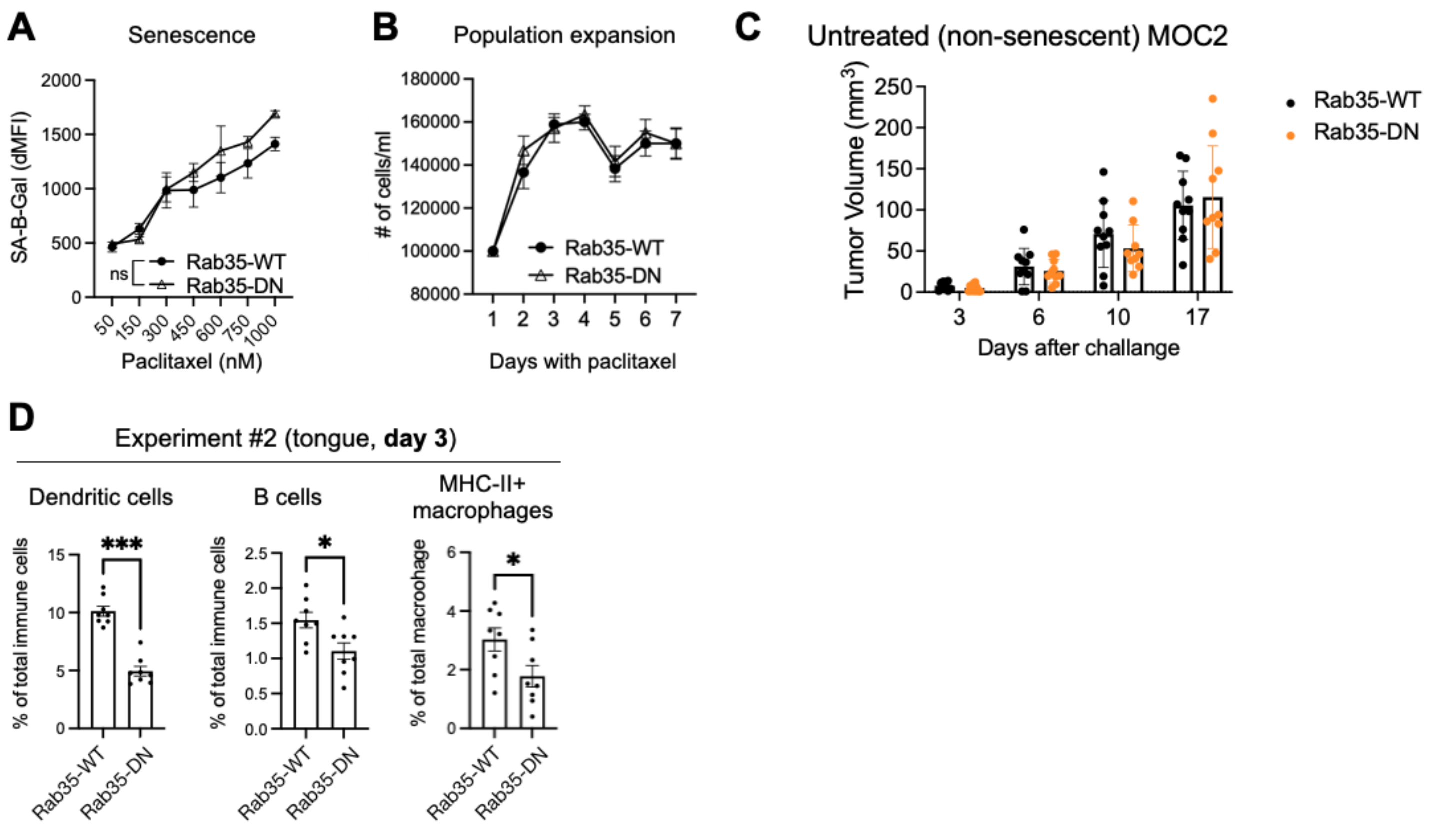


### Figure S4. Inhibition of tumor recurrence is a specific property of senEVs

**(A-B)** Analysis of SA-B-Gal activity (A) and cell expansion (B) in Rab35-WT and Rab35-DN senescent MOC2 cells cultured *in vitro* demonstrates that Rab35 manipulation does not influence senescence induction, as there is no significant difference in the senescence-inducing effect of paclitaxel. Statistical tests by one-way ANOVA. **(C)** Tumor growth of untreated MOC2 cells expressing Rab35-WT or Rab35-DN indicate that MOC2 tEVs do not impact tumor progression like senEVs do. **(D)** Quantification of immune cell infiltrates by flow cytometry 3 days after implantation of Rab35-WT or Rab35-DN senescent MOC2 cells (related to Figure 4D). Statistical tests by Mann-Whitney.

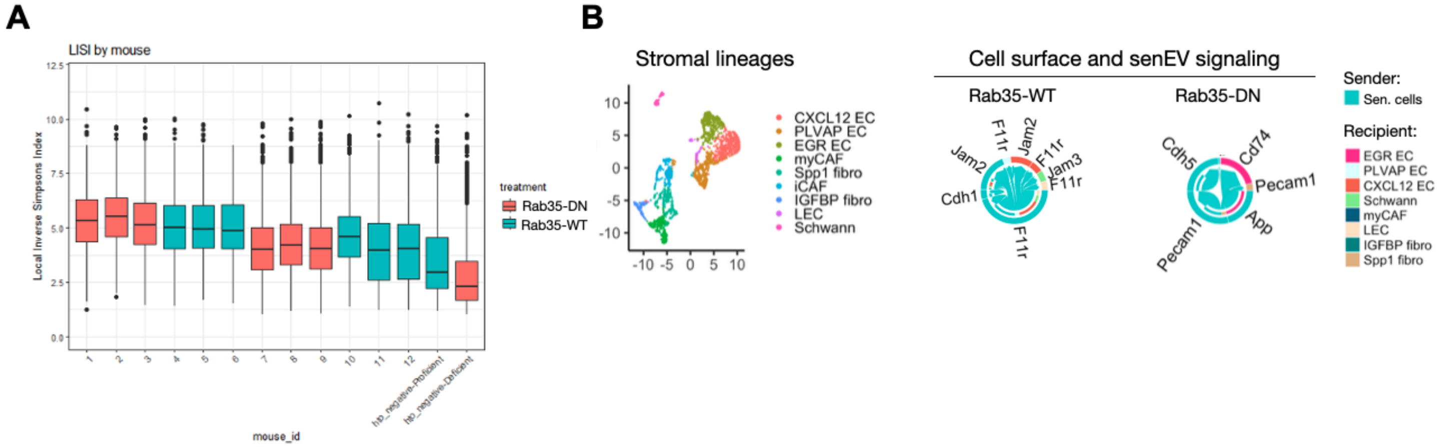


Figure S5. Senescent cells are capable of binding to endothelial cells but not to fibroblasts

**(A)** After SCTransform (to correct for the read-depth disparity between two independent single cell sequencing experiments), the Local Inverse Simpson's Index (LISI) shows that the samples were well mixed and lacked any evidence of further batch effects requiring correction. **(B)** Sub-clustering of cells belonging to the stromal lineage based on differentially expressed genes compared to other lineages (left). Visualizing cell-cell communications mediated by membrane-bound ligand-receptor interactions using chord diagrams (right). Signaling between senescent cells and stromal cells is shown. Each sector in the chord diagrams is an arrow depicting the direction of the ligand-receptor signaling.

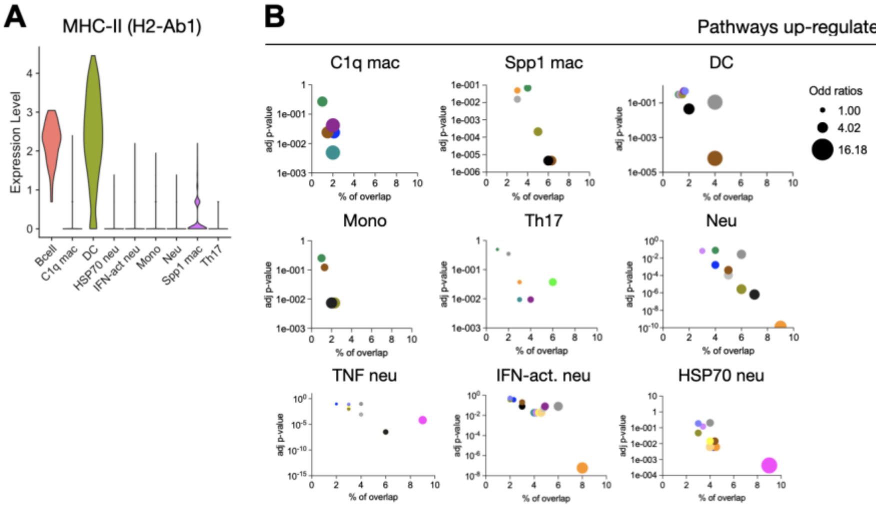


| | C1q mac | Spp1 mac | Mono | Mo-DC | Neu | TNF neu | IFN-act. neu | HSP70 neu | Th17 |
|---|---|---|---|---|---|---|---|---|---|
| Angiogenesis | | | | | | X | | X | |
| Apoptosis | | | | | | | X | X | |
| Complement | X | X | X | X | X | | X | X | |
| EMT | | X | X | X | X | X | X | X | |
| IFN-a response | X | | | X | | | X | | X |
| IFN-g response | X | | | X | | | X | | X |
| IL-2/STAT5 Sig. | X | | | | X | X | X | X | |
| IL-6/JAK/STAT3 Sig. | X | X | X | | X | | | | X |
| KRAS Sig. | | X | X | X | X | X | X | X | |
| mTORC1 Sig. | | X | | X | X | X | X | X | |
| Myc Targets V1 | | | | X | X | | X | X | |
| Notch Sig. | | | | | | | | | X |
| TGF-beta Sig. | | | | X | X | X | X | X | X |
| TNF-a via NF-kB | | X | | | X | X | X | X | X |
| p53 Path. | | | | | | X | X | X | |
| P13K/AKT/mTOR | | | | X | | X | X | X | |

Figure S6. Pathways up-regulated in immune cells in presence of senEV-proficient senescent cells

**(A)** Expression of the MHC-II gene (H2-Ab1) by immune cells identifies *bona fide* antigen presenting cells. **(B)** Genes significantly up-regulated in immune cells define signaling pathway active in EV-proficient senescent microenvironments (as compared to EV-deficient senescent microenvironments).

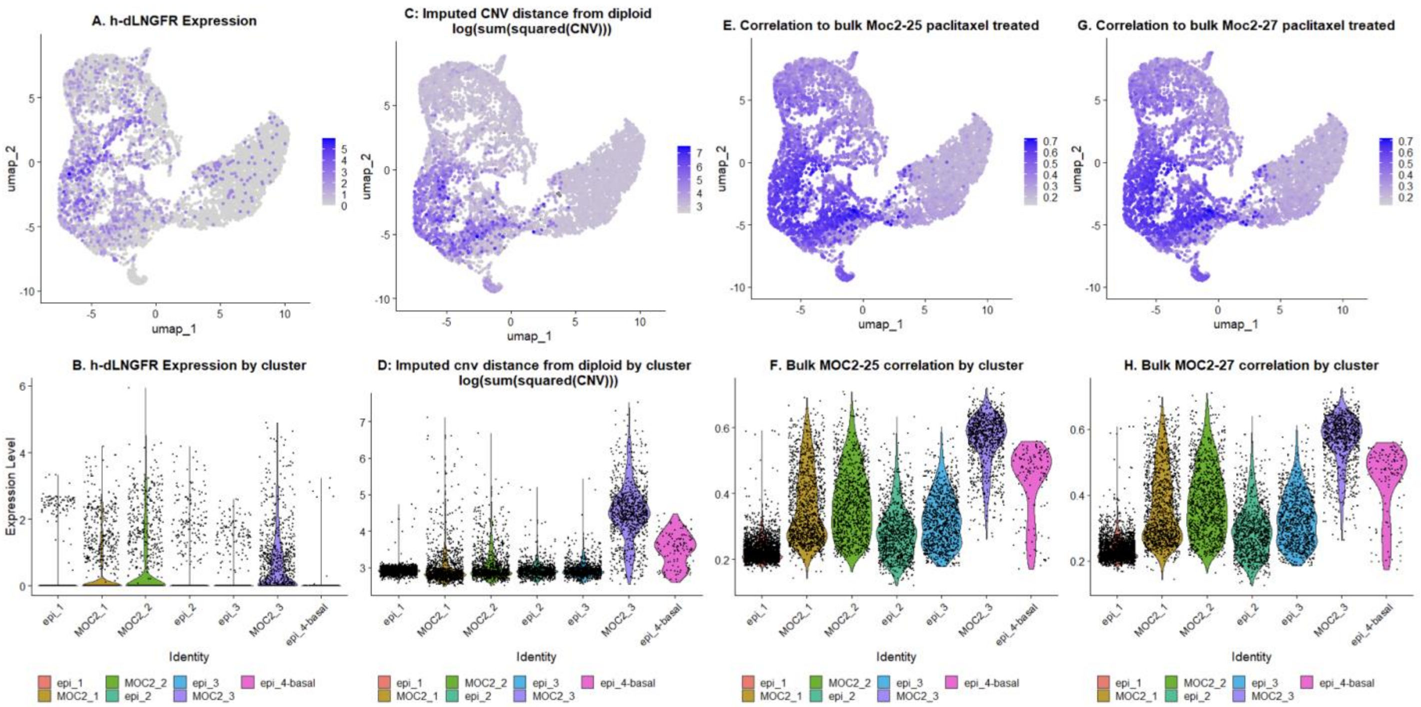


#### Figure S7. Implanted MOC2 cells have high h-dLNGFR expression, copy number distance from diploid and correlation to bulk MOC2 expression

**(A, C, E, G)** Feature plots showing the UMAP embedding of the epithelial cell subset colored by h-dLNGFR expression (A), imputed copy number distance from diploid (C)**,** correlation to Rab35-WT MOC2 (MOC2-25) bulk sequencing, and (G) correlation to Rab35-DN MOC2 (MOC2-27) bulk sequencing. **(B, D, F, H)** Violin plots showing the single-cell values of features plotted in (A, C, E, G).

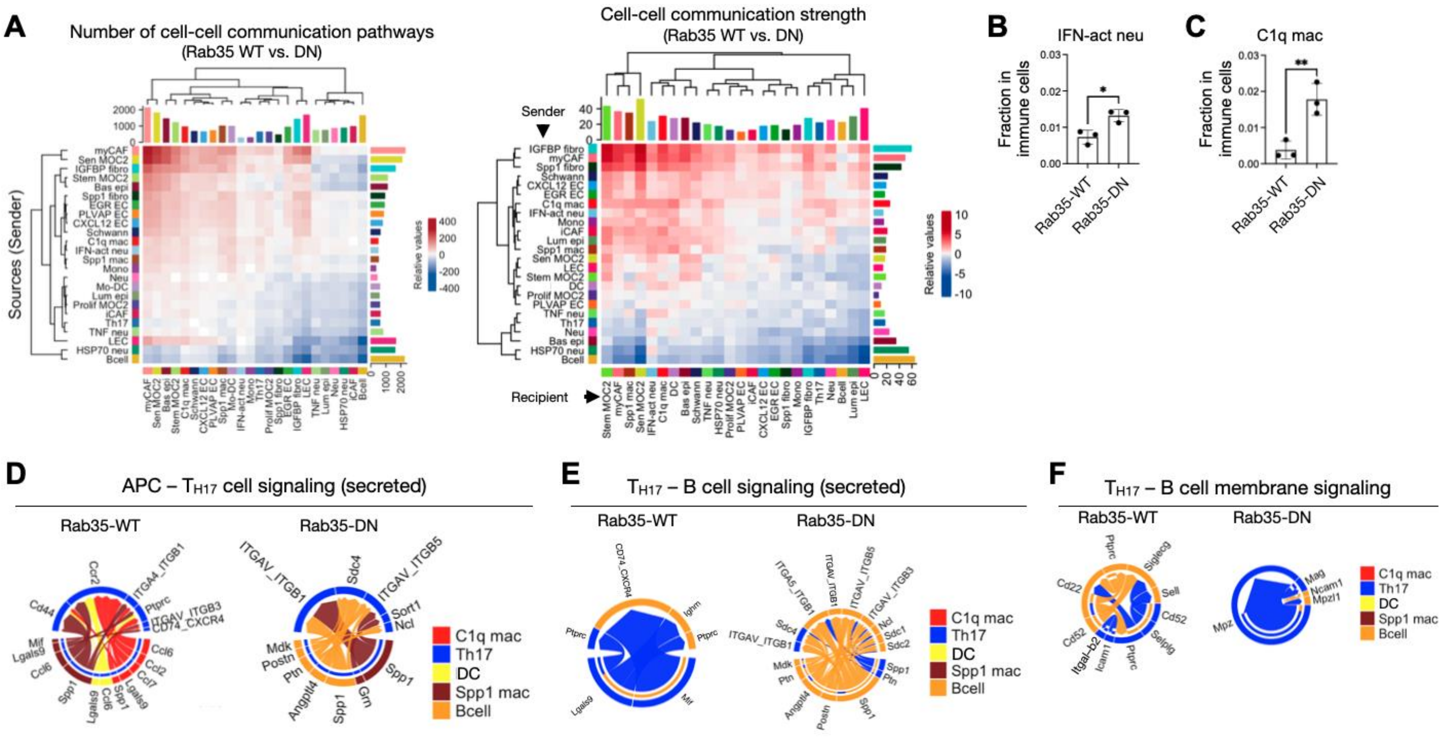


Figure S8. The presence of senEVs in the senescence microenvironment impacts downstream immune responses

**(A)** Analysis of cell-cell interaction numbers (left) and cell-cell interaction strength (right) within EV-proficient senescence microenvironment (compared to EV-deficient). The number/strength of ligand-receptor interactions among the 24 identified cell types is color-coded as red (increased) or blue (decreased). The bar plots on top and right-side represent the sum of each column and row, respectively, of the absolute values displayed in the heatmap (incoming and outgoing signaling, respectively). **(B-C)** Increase in the frequency of IFN-activated neutrophils (B) and C1q macrophages (C) in senEV-deficient senescence microenvironments. **(D-F)** Visualizing cell-cell communications mediated by secreted (D-E) or membrane (F) ligand-receptor interactions using chord diagrams. Signaling between antigen-presenting cells (dendritic cells, B cells and spp1 macrophages) plus C1q macrophages and Th17 cells (D), and B cells and Th17 cells (E-F) is shown. Each sector in the chord diagrams is an arrow depicting the direction of the ligand-receptor signaling.

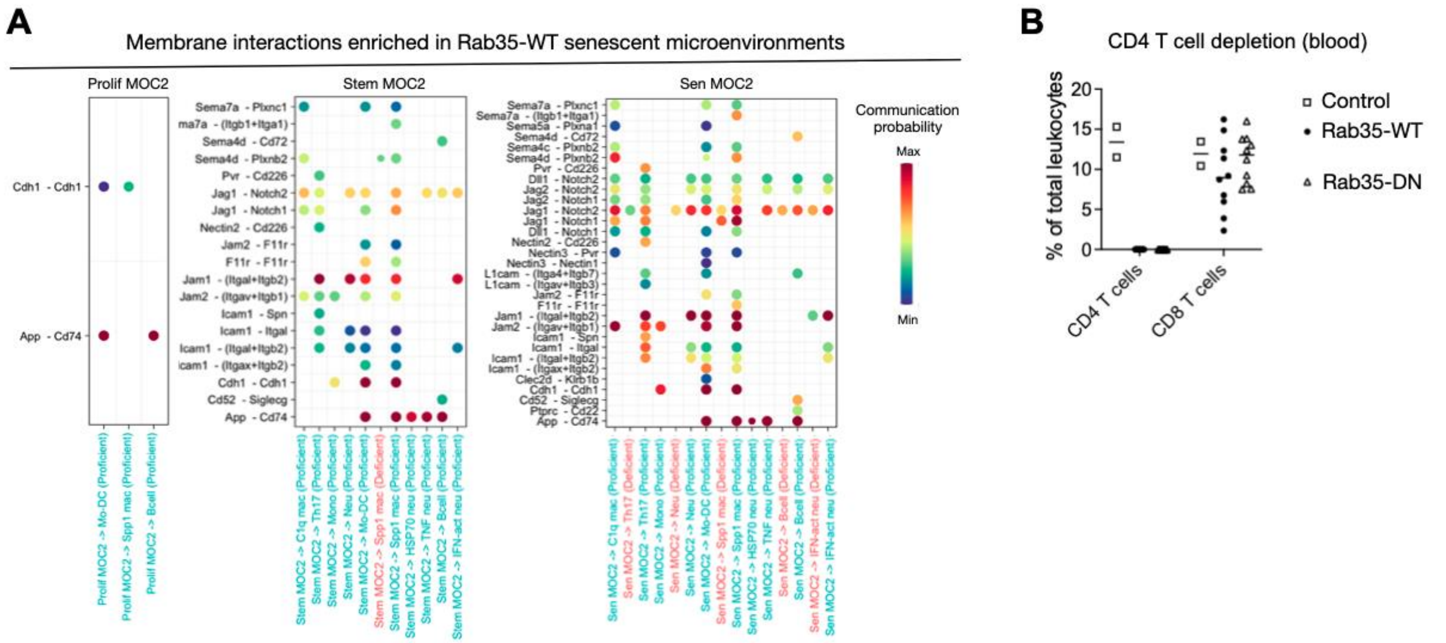


Figure S9. Membrane-bound ligand-receptor interactions between senescent and immune cells overlap among senescent cell subsets.

**(A)** Bubble plots showing enriched interactions between senescent and immune cells in presence of senEVs. Each senescent cell subset is shown separately. The ligand-receptor pair may be found either exclusively in the context of EV-proficient senescence microenvironments, or in both EV-proficient and EV-deficient senescence microenvironments but upregulated only in EV-proficient cells. The color bar represents the communication probability (interaction strength) in the second dataset relative to the first dataset. **(B)** Flow cytometric quantification of T cell subsets in blood of mice treated with CD4 depleting antibodies 4 days before (related to Figure 5H). Control mice are untreated.
